# Secretin receptor as a target in gastrointestinal cancer: expression analysis and ligand development

**DOI:** 10.1101/2022.01.31.478443

**Authors:** Anja Klussmeier, Stefan Aurich, Lars Niederstadt, Bertram Wiedenmann, Carsten Grötzinger

## Abstract

Secretin was originally discovered as a gastrointestinal peptide that stimulates fluid secretion from pancreas and liver and delays gastric emptying. In disease, secretin receptor (SCTR) was found to occur as a splice variant in gastrinoma and pancreatic adenocarcinoma. Overexpression of SCTR has been described for gastrinomas, carcinoid tumors of the lung and cholangiocarcinoma. SCTR therefore remains a candidate target for molecular tumor imaging as well as for peptide receptor radioligand therapy (PRRT) in a number of oncological indications. The aim of this study was to characterize SCTR expression in esophageal and pancreatic cancer. 65 of 70 PDAC tissues stained strongly positive for SCTR in immunohistochemistry, as were most of 151 esophageal cancer samples, with minor influence of grading in both entities. In addition, the aim of this study was to further delineate residues in human secretin that are critical for binding to and activation of the human SCTR. For a potential development of short and metabolically stable analogs for clinical use, it was intended to probe the peptide for its capacity to incorporate deletions and substitutions without losing its affinity to SCTR. A library of 146 secretin variants containing single amino acid substitutions as well as truncations on either end was tested using β-arrestin2-GFP translocation and fluorescent ligand internalization high-content analysis and cAMP assays run in agonist and antagonist mode as well as a radioligand binding. The main structural determinants of SCTR binding and activation were localized to the N-terminus, with His^1^, Asp^3^ being among the most sensitive positions, followed by Phe^6^, Thr^7^ and Leu^10^. Aminoterminal truncation caused a rapid decline in receptor activity and most of these variants proved to be partial agonists showing antagonistic properties. In this study, the most potent antagonist showed an IC_50_ of 309 ± 74 nM in the β-arrestin2-GFP translocation assay on human SCTR while remaining a weak partial agonist. Future studies will have to demonstrate the utility of further enhanced secretin analogues as tracers for in vivo imaging and therapy.

## 1. Introduction

Secretin is a peptide hormone produced in the S cells of the small intestinal mucosa [1]. It was originally discovered as a gastrointestinal peptide that stimulates fluid secretion from pancreas [2] and liver [3] and delays gastric emptying [4]. More recently, it has been implicated in the regulation of food intake [5], drinking behavior and water homeostasis [6–8], as well as motor behavior, learning and memory [9]. A 121 amino acid precursor protein, preprosecretin, is the initial product of transcription and translation of the secretin gene. During the maturation process, the proteolytic cleavage of a signal peptide and associated peptides finally releases the 27 amino acid form of secretin [9,10]. While its helical part seems to mediate high-affinity binding to the receptor, the unstructured domain has long been known to be responsible for activation of the secretin receptor (SCTR) [11–13]. A high homology of the N-terminal part of the peptide ligands of other class B GPCRs has been identified [14–16]. Several studies have contributed to a better understanding of the role of secretin primary and secondary structure for receptor binding and activation [17–20]. Recently, the three-dimensional structure of a secretin-SCTR-Gs complex has been resolved using cryogenic electron microscopy, revealing a unique spatial organization of the SCTR extracellular domain relative to the heptahelical membrane core [21].

Alternative cleavage during maturation can lead to secretin variants [22]. In the maturation process of porcine secretin, e.g., two alternative peptides with unknown function have been identified, elongated at the C-terminus either by Gly or Gly-Lys-Arg. In vivo, these naturally occurring variants display an activity comparable to native secretin [23]. Experiments utilizing N-terminally truncated variants (amino acids 2-27) displayed a severe reduction in activity on isolated pancreatic membranes, while the truncation to amino acids 7-27 completely abolished agonistic effects [24]. Despite their lack of activation, these truncated peptides were able to competitively displace native secretin at high concentrations [24,25]. In lack of more potent antagonistic analogues, until the recent creation of a high-affinity SCTR antagonist [26], they continued to be utilized as antagonists in experimental research.

The secretin receptor (SCTR), initially cloned in 1991, represents the archetype of the B family of GPCRs [27–30]. This receptor family early attracted the interest of pharmaceutical research and development, since it contains a variety of important targets like the receptors for VIP, GLP-1, GLP-2, PACAP and calcitonin [15,30,31]. Upon activation by its ligand, SCTR signals through the increase of intracellular cAMP via Gs and by stimulation of calcium mobilization via Gq proteins [12,32,33]. In parallel, receptor and ligand are internalized to allow receptor turnover or degradation [34]. In cancer, SCTR was found to occur in a misspliced form in gastrinoma and pancreatic adenocarcinoma [35–37]. Whether this splice variant acts in a dominant-negative manner in tumors, remains a matter of debate [38]. Overexpression of SCTR has been described e.g., for gastrinomas, carcinoid tumors of the lung and cholangiocarcinoma, but was not found in hepatocellular carcinoma [39–41].

SCTR thus constitutes a candidate target for molecular imaging as well as for peptide receptor radioligand therapy (PRRT) in a number of oncological indications. Somatostatin receptors have proven to be instrumental in targeting neuroendocrine neoplasms for the past two decades [42,43] and similar approaches are warranted for tumors with a poor prognosis such as pancreatic and esophageal cancer. Therefore, this study aimed at evaluating SCTR expression in cancers originating from esophagus and pancreas. For a potential development of truncated and metabolically stable analogs for clinical use, this study was designed to probe the secretin peptide for its capacity to incorporate deletions and substitutions without losing its activity and affinity. Consequently, the structure-activity relationship of secretin was investigated by analyzing a library of variants of the natural peptide, with the intent of paving the way for the future development of secretin-based tumor tracers.

## 2. Materials and Methods

### 2.1. Cell lines

U2OS-SCTR/β-arr2-GFP: human osteosarcoma cell line U2OS) purchased from ATCC/LGC Standards, Wesel, Germany) were stably transfected with plasmids encoding SCTR (*Homo sapiens*) and the fusion protein β-arrestin2-green fluorescent protein (GFP) (*Rattus norvegicus-Aequorea victoria*) (β-arr2-GFP).

### 2.2. Buffers, reagents and peptides

If not indicated otherwise, all chemicals were purchased from Carl Roth GmbH, Karlsruhe, Germany. Assay buffer C1: 130 mM NaCl; 5 mM KCl, 10 mM HEPES, 2 mM CaCl_2_, 10 mM glucose, 1 % BSA. Membrane isolation buffer: 5 mM Tris-HCl pH 7.6, 5 mM MgCl_2_, 1 mM EGTA, cOmplete protease inhibitor cocktail (Roche, Mannheim, Germany). Membrane binding buffer: 50 mM HEPES pH 7.4, 1 mM CaCl_2_, 2.5 mM MgCl_2_, 0.5 % BSA, cOmplete protease inhibitor cocktail. Membrane Wash buffer: 50 mM Tris-HCl pH 7.4, 125 mM NaCl, 0.05 % BSA. Peptides were supplied by peptides&elephants (Hennigsdorf, Germany).

### 2.3. Software

For dose-response evaluation, Spotfire Decision Site (TIBCO, Palo Alto, CA, USA) was used to calculate a four-parameter logistic regression curve fit [y=min+(max-min)/(1+(EC_50_/x)^Hill^)].

### 2.4. Immunohistochemistry

For the characterization of SCTR in normal and pathological human tissues, commercially available tissue microarrays (TMAs) were employed (Accumax A307 and A218 [Biocat, Heidelberg, Germany], PA2081 and BS02051 [Biomax, Rockville, MD, USA], BR7 and CR1 [SuperBioChips, Seoul, Republic of Korea]. Paraffin-embedded TMAs were deparaffinized and cooked in 10 mM citrate buffer (pH 6,0) for 4 min. Sections were then washed in PBS and blocked for 5 min in 3% H2O2, and for an additional 30 min in 5% goat serum in PBS. After washing, sections were incubated over night at 4°C with an anti-SCTR antibody (HPA007269, Sigma-Aldrich) (0.4 μg/ml). The following day, the sections were PBS-washed again and incubated with a biotinylated goat anti-rabbit antibody for 30 min at room temperature followed by a 30 min incubation with AB-complex (VectorLabs, Burlingame, CA, USA). The brown staining was then generated by incubation of the sections with Liquid DAB+ Substrate Chromogene Kit (Dako, Glostrup, Denmark) for 2 min. Finally, tissue sections were counterstained for 10 min with Mayer’s Hämalaun solution (SigmaAldrich). Sections were then mounted, dried, and representative pictures were taken with an Axio Observer.Z1 microscope (Zeiss, Jena, Germany). Two methods of quantitative estimation of SCTR staining were applied. For esophageal normal and tumor tissues, the extent E (percentage of area stained, ranging from 0% to 100%) was determined by an algorithm using Axiovision software (Zeiss), while intensity of the staining (I) was estimated by two observers using a range from 0 to 3. The immunohistochemistry score (IHC score) was calculated by multiplying values for extent and intensity (IHC score = E*I), yielding numbers between 0 and 300. For pancreatic normal and tumor tissues, SCTR detection was estimated in a single parameter (strength of staining) with three discrete values as either strong staining, weak staining or no staining.

### 2.5. Agonist-induced β-arr2-GFP translocation

To evaluate reactivity of human SCTR exposed to different ligand variants, intracellular translocation of a fusion protein of β-arrestin2 and green fluorescent protein was utilized. For evaluation of activation, receptor-mediated recruitment of cytoplasmic β-arr2-GFP was monitored through its translocation into condensed intracellular vesicles. The extent of translocation events was quantified by determining the increasing granularization of the intracellular fluorescence utilizing an automated high-throughput microscope in combination with computational algorithms to estimate potency (EC_50_) and efficacy, thus characterizing ligand-SCTR interaction (Figure 3). U2OS-SCTR/β-arr2-GFP cells were seeded in 96-well cell culture plates to 60-80 % confluence. On the following day, preceding stimulation, cells were incubated for one hour in serum-free medium. Ligand variants were applied as serial dilutions in assay buffer C1. After preincubation, supernatant was discarded by tapping plates overhead repeatedly. For stimulation, 50 μl of ligand dilution was applied per well in duplicates and incubated for 20 minutes at 37 °C. Successively, cells were fixed by applying 4 % buffered formaldehyde (Herbeta Arzneimittel, Berlin, Germany) for 10 minutes at room temperature and consecutively permeabilized for 10 min at room temperature with 0.1 % TritonX-100 in PBS, containing DAPI (4’,6-Diamidin-2-phenylindol, SigmaAldrich, Deisenhofen, Germany, 1 μg/ml) to counterstain nuclei. Microscopic images were recorded by an IN Cell Analyzer 1000 (GE Healthcare, Reading, UK) with 20x magnification at 5 frames per well in a 96-well format. Image processing was performed with IN Cell Investigator software (GE Healthcare), applying the provided granularity algorithm. The output of the analysis was indicated as vesicle area/cell and was utilized for calculation of concentration-response curves. Parameters derived from curve fitting included the concentration of half-maximal activity (EC_50_), and intrinsic activity. Intrinsic activity was normalized defining reaction to non-modified wild type secretin as the 100 % curve maximum value. Values displayed in this study represent the mean of at least three independent experiments, independently performed in duplicates.

### 2.6. Evaluation of antagonistic function/IDCC internalization assay

For analysis of antagonistic effects on SCTR, a slightly modified protocol of the β-arr2-GFP translocation was used. Ligands with suspected antagonist function were incubated with cells as previously described for 5 min at 37 °C, followed by the addition and incubation with secretin covalently linked to an indodicarbocyanine (IDCC) fluorophore (secretin-IDCC, 10 nM, 15 min at 37 °C). For analysis, cells were terminally fixed and permeabilized as previously described. The additional monitoring of IDCC fluorescence in the near-infrared channel (667 nm) provided another measure for receptor binding and activation.

### 2.7. Measurement of intracellular cAMP concentrations in transfected cell lines

Analysis of downstream signaling of activated SCTR via Gαs G-protein subunits was performed by monitoring intracellular cAMP utilizing the Promega GloSensor system in accordance with the supplier protocol. Measurements were performed using the EnVision Analyzer (PerkinElmer, Rodgau, Germany). For evaluation, concentration-response curves (signal-to-noise ratios plotted) employing the derived statistical mean and SD for estimation of the respective EC_50_ were calculated using Spotfire Decision Site (TIBCO). For the analysis of antagonists, a modified protocol was used. Treatment of cells and dilution of the potential antagonist was performed as previously described. Ligand dilutions were added to cells and incubated at room temperature for 5 min. Successively, secretin was added at a final concentration of 0.5 nM for an additional 15 min at room temperature. The emitted chemiluminescence, which correlates with the cAMP concentration, was detected with suitable filters. Evaluation of data was performed as previously described with a calculation of half-maximal inhibitory concentration (IC_50_).

### 2.8. Radiolabeling of secretin

For the evaluation of ligand binding to SCTR, native human secretin was slightly modified from its original amino acid sequence. To increase spectroscopic absorption during HPLC purification of the radiolabeled peptide, residue 27 was changed from valine to tryptophan. To allow the covalent linkage of ^125^I to an aromatic side chain, an additional tyrosine was introduced at the C-terminal end of the peptide (W27, Y28-secretin). Radiolabeling was then performed on the modified peptide through an adaption of the chloramin-T method initially introduced by Hunter and Greenwood [44]. To achieve an effective reaction, the peptide was diluted in 0.5 M sodium phosphate buffer (pH 7.6) to a concentration of 200 μM with an addition of 0.5 mCi carrier-free ^125^I-NaI in 2 μl of carrier solution (PerkinElmer). Iodination was initiated by addition of 4 μl chloramin-T solution (SigmaAldrich, 1 mg/ml). The reaction was terminated by addition of 4 μl of sodium metabisulfite (2 mg/ml). Analysis and separation of resulting radiopeptide fractions was carried out via HPLC. The separation after injection of the complete iodination reaction was performed utilizing a Zorbax 300SB-C18 column (Agilent, Waldbronn, Germany) with elution of bound peptide by a water-acetonitrile gradient from 10 to 60% containing 0.1% TFA within a runtime of 60 minutes.

### 2.9. Isolation of SCTR-bearing membrane fractions

U2OS cells overexpressing human SCTR were cultivated to confluence and washed with PBS once. To detach the cultured layer, cells were moistened with ice cold PBS containing 5 mM EGTA, scraped into suspension and centrifuged at 4 °C/ 200 g for 10 minutes. Supernatants were discarded and cell pellets were resuspended in membrane isolation buffer. Homogenization was achieved by applying shearing force using a precooled Dounce homogenizer on ice with subsequent sedimentation of membrane fractions by ultracentrifugation at 4° C at 40.000 x g for 45 minutes. Pellets were resuspended in isolation buffer, homogenized and centrifuged again as described. The final purified membrane pellet was carefully resuspended in membrane binding buffer without supplementation of BSA, snap-frozen in liquid nitrogen and stored at −80° C until used.

### 2.10. Radioligand binding assay

10 μg of membrane extracts from SCTR-overexpressing U2OS cells were resuspended in membrane binding buffer containing 0.5 % BSA and supplemented with 160 pM of I^125^ labeled secretin (quantification according to specific I^125^ activity assuming monoiodination). Depending on the expected range, unlabeled competing ligands were diluted in membrane binding buffer at a final concentration between 0.1 fM and 10 μM. To allow binding, suspensions were incubated for 1 h at 37 °C and consecutively transferred to filter plates precoated with 0.25 % polyethylenimine (average molecular weight ~60.000, SigmaAldrich) in H2O. After capturing membrane fractions and bound ligands through suction, the filter material was washed six times with membrane wash buffer and dried at 50 °C for 15 minutes. Evaluation of bound radioligand was performed by scintillation count utilizing UltimaGold substrate solution (PerkinElmer) and a MicroBetaII plate-based scintillation counter (PerkinElmer). Scintillation readout was used to calculate concentration-response curves and IC_50_ values. Values were normalized over mean plateau of maximum scintillation intensity values defined as 100 and used for composition of concentration-response curves.

## 3 Results

### 3.1 SCTR Expression in human tissues

To evaluate the role of SCTR as a potential target in tumors of the gastroenteropancreatic duct, paraffin sections of normal and cancerous tissue samples were to be examined using immunohistochemistry (IHC). In order to obtain robust and reliable results, a commercially available SCTR antibody was validated and found to show the expected plasma membrane staining as well as a high specificity for SCTR (Supplementary figure 1). Using this antibody, 181 samples from normal esophagus and esophageal cancer were analyzed and the obtained microscopic images were used to score SCTR staining extent and intensity. In normal esophagus, the IHC score was found to be low (23.9 ± 16.3, mean ± SD) and to be restricted to the basal cell layer of the mucosa (Figure 1). In all analyzed cancer types, the SCTR antibody showed mostly intense staining in almost all tumor cells, with the IHC score being much higher: 185.8 ± 86.3 for esophageal squamous cell carcinoma (ESCC), 100.7 ± 79.8 for esophageal adenocarcinoma (EAC) and 145.8 ± 33.2 for esophageal adenosquamous carcinoma (EASC) (Figure 1).

**Figure 1.**
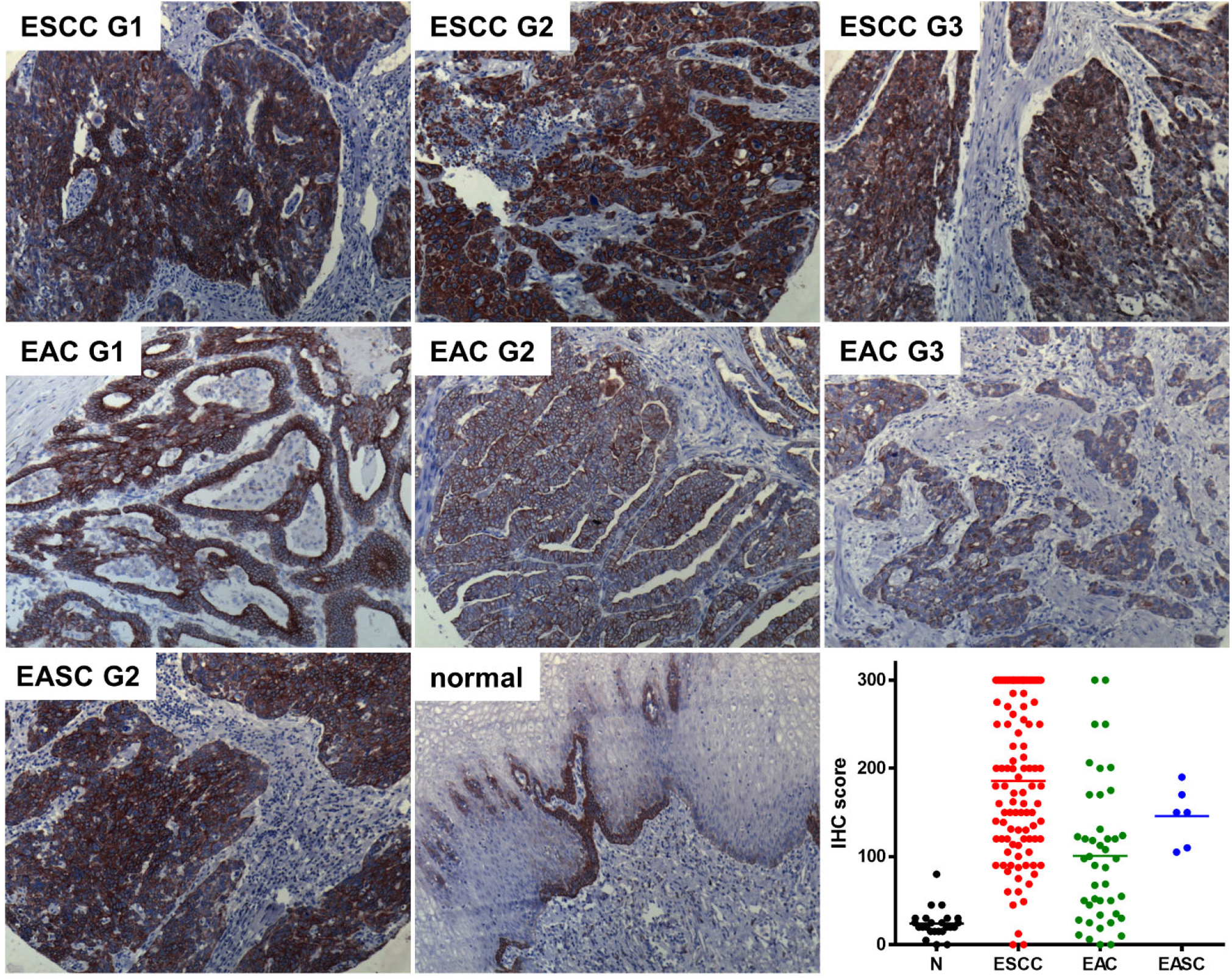
Immunohistochemical detection and quantification of SCTR in esophageal cancer of various histopathological grades (G1, G2, G3) and in normal human esophagus. Representative SCTR immunohistochemistry stainings are presented for ESCC, EAC, EASC and non-cancerous esophagus tissue (normal). In addition, a quantification of the staining (IHC score) for n=30 normal tissue samples (N) as well as for the three tumor entities ESCC (n=101), EAC (n=44) and EASC (n=6) is presented. Microscopic images were taken using a 10x objective.

With regard to histopathological grade, SCTR staining was found highest in well-differentiated G1 ESCC and EAC, yet IHC score was only gradually lower in G2 and typically still high even in G3 tumors as compared to normal esophageal tissue (Figure 1 and Supplementary figure 2).

In addition to esophageal cancer, expression of SCTR was evaluated in pancreatic ductal adenocarcinoma (PDAC). In normal pancreas, ductal epithelial cells were highlighted by SCTR antibody staining, while pancreatic acinar cells and cells in the islets of Langerhans were not stained. Besides normal pancreas, 70 PDAC tissues as well as samples of 3 adenosquamous carcinomas, one malignant islet cell carcinoma, 10 benign islet cell tumors, two cases of ductal hyperplasia and 10 cases of chronic pancreatitis were examined. While non-tumor cells were not stained, the majority of tumor cells in the cancerous areas were SCTR-positive. 93% of all PDACs were found to show strong SCTR staining, including 19 out of 23 (83%) of poorly differentiated G3 PDAC tissues (Figure 2, Table 1). All well-differentiated (G1) PDACs (n=10) and 36 out of 37 G2 PDACs (97%) stained strongly positive for SCTR. In all cases of chronic pancreatitis (n=10), SCTR staining was no more pronounced than in normal pancreas.

**Figure 2.**
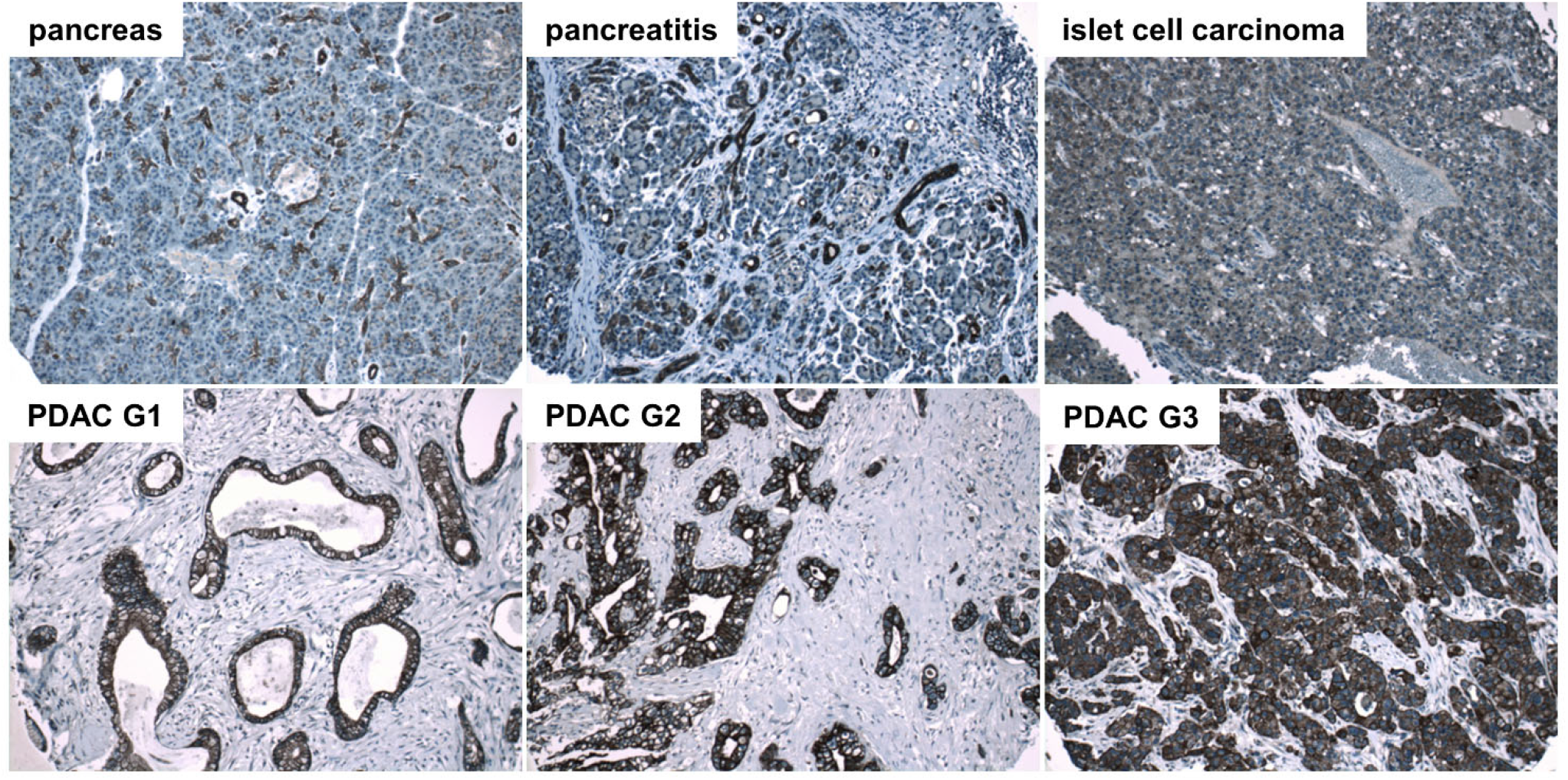
Immunohistochemical detection of SCTR in human normal pancreas, chronic pancreatitis and islet cell carcinoma as well as in PDAC of histopathological grades G1, G2 and G3. Representative SCTR immunohistochemistry microphotographs are presented. Microscopic images were taken using a 10x objective.

**Table 1.** Quantification of SCTR staining intensities scored as strong, weak or absent in PDAC and other pancreatic pathologies.

| Tissue | Grade | Strong<br>SCTR staining | Weak<br>SCTR staining | No<br>SCTR staining |
| --- | --- | --- | --- | --- |
| PDAC | All | 65/70 (93 %) | 1/70 (1 %) | 4/70 (6 %) |
| PDAC | G1 | 10/10 (100 %) | 0/10 (0 %) | 0/10 (0 %) |
| PDAC | G2 | 36/37 (97 %) | 1/37 (3 %) | 0/37 (0 %) |
| PDAC | G3 | 19/23 (83 %) | 0/23 (0 %) | 4/23 (17 %) |
| Adenosquamous carcinoma |  | 2/3 (67 %) | 1/3 (33 %) | 0/3 (0 %) |
| Islet cell carcinoma (malignant) |  | 0/1 (0 %) | 0/1 (0 %) | 1/1 (100 %) |
| Islet cell tumor (benign) |  | 0/10 (0 %) | 1/10 (10 %) | 9/10 (90 %) |
| Ductal hyperplasia |  | 2/2 (100 %) | 0/2 (0 %) | 0/2 (0 %) |
| Chronic pancreatitis |  | 0/10 (0 %) | 0/10 (0 %) | 10/10 (100 %) |

In addition to samples from esophagus and pancreas, a number of normal samples from other tissues were examined for SCTR expression, including stomach, intestine, liver, lung and kidney. While brain, spleen, lymph node, lung and adrenal gland were found to be SCTR-negative, most tissues with ductal or other epithelia stained clearly positive, including stomach, small intestine, rectum, gallbladder and common bile duct. In the liver, only bile ducts stained positive and in kidney, distinct structures, most likely of the ascending loop of Henle or collecting tubules were SCTR-positive (Supplementary figure 3).

### 3.2 Assay development

To determine the activity of a large number of different secretin variants, a robust and flexible high-throughput assay had to be established. The transfluor technology was chosen, as it provides for a simple, reliable and high-throughput-compatible readout system with favorable signal-to-background ratios [45,46]. U2OS cells stably co-expressing a β-arrestin2-GFP fusion protein (β-arr2-GFP) and human SCTR were used to set up a high-content analysis system (automated microscopy and image analysis) that would allow characterizing SCTR activation by a large number of ligands. This system was designed to accommodate assays run in agonist as well as in antagonist mode. Fig. 3A shows the morphological transition of β-arr2-GFP from a diffuse distribution throughout the cytoplasm to a pronounced vesicular pattern found after incubation with 10 nM secretin for 30 min. The segmentation algorithm developed for image analysis provided for an efficient quantification of vesicle areas. These were plotted versus peptide concentration and allowed derivation of concentration-response curves and the calculation of potency (EC_50_) as well as efficacy (Fig. 3B).

**Figure 3:**
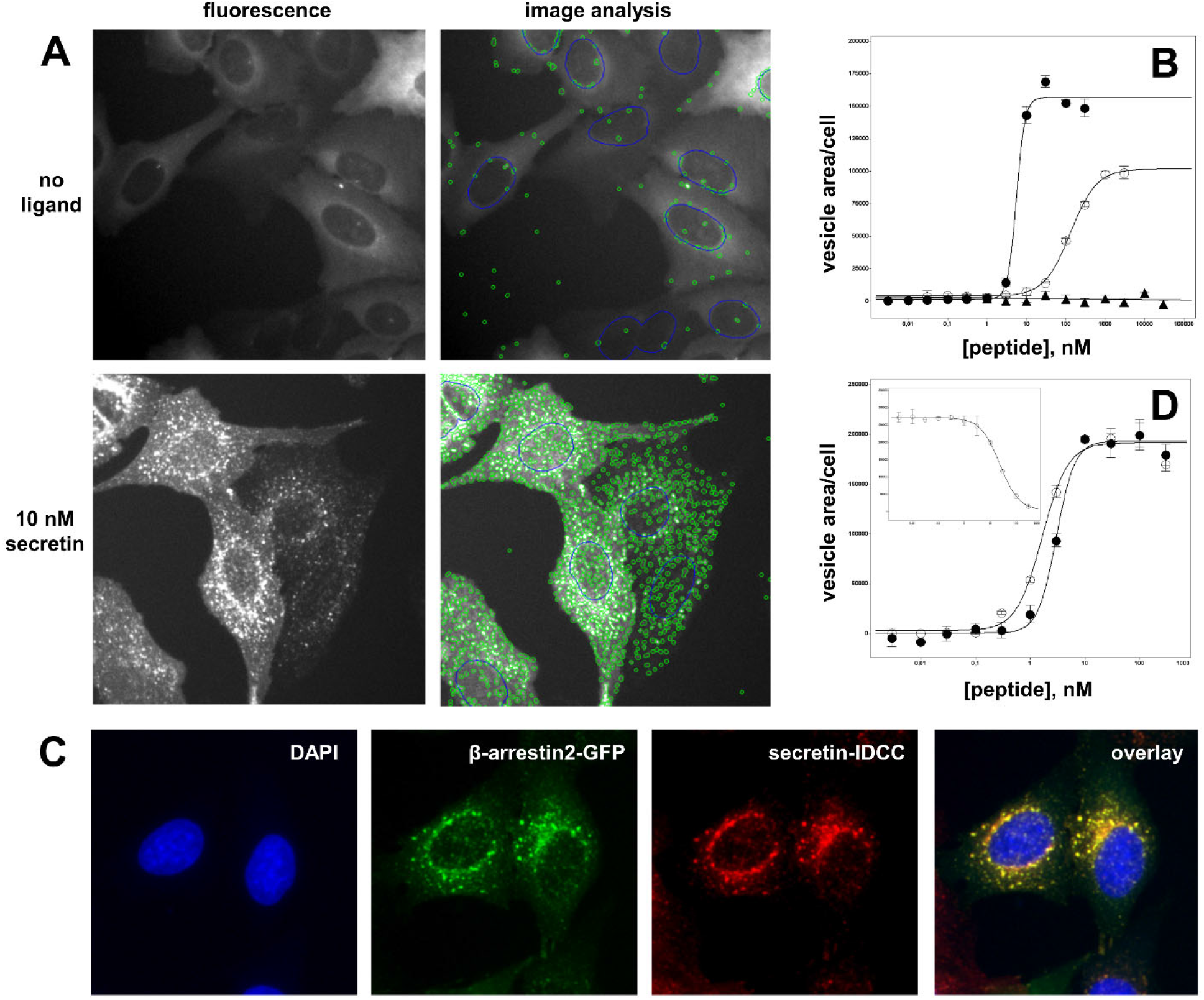
High-throughput quantification of receptor activation through measurement of β-arrestin translocation. **A)** U2OS cells co-expressing β-arr2-GFP and SCTR were utilized to quantify SCTR activation through automated microscopy and image analysis. In the resting state, without ligand application, β-arr2-GFP localizes diffusely to the cytoplasm, only few vesicles were detected (upper row). After 20 min of incubation with 10 nM secretin, β-arr2-GFP localizes to a multitude of bright vesicles that were recognized and counted by the segmentation algorithm (lower row). **B)** Quantitative data from image analysis were plotted in concentration-response curves and derivation of potency (EC_50_) and intrinsic activity values for SCTR agonism. Closed circles (●): wild-type secretin (EC_50_ = 3.26 ± 0.80 nM), open circles (○): Ala^6^-secretin (EC_50_ = 155.2 ± 30.8 nM, efficacy = 61.8 ± 4.9%), closed triangles (▲): wild-type GLP-2 (no SCTR activation). **C)** Microscopic representation of the ligand-induced translocation of β-arr2-GFP (green) and internalization of fluorescently labelled secretin-IDCC (red), nuclear staining (DAPI) shown in blue. The yellow color in the overlay indicates vesicular co-localization of fluorescently labelled ligand and β-arr2-GFP. **D)** Quantification of internalized secretin (secretin-IDCC) showed equal potency and efficacy values as the β-arr2-GFP translocation assay: closed circles (●): secretin-IDCC, quantified in the β-arr2-GFP translocation assay (EC_50_ = 3.02 ± 0.12 nM), open circles (O): secretin-IDCC, quantified in secretin-IDCC internalization assay (EC_50_ = 1.45 ± 0.54 nM). Inset: secretin-IDCC internalization assay run in antagonist mode (displaced by secretin).

Wild-type secretin showed an EC_50_ of 3.26 ± 0.80 nM and an intrinsic activity of 100% in this assay, while Ala^6^-secretin was found to have an EC_50_ of 155.2 nM ± 30.8 nM and an intrinsic activity of 61.8 ± 4.9%, while the structurally related glucagon-like peptide 2 (GLP-2) showed no SCTR activation (Fig. 3B). As SCTR activation is accompanied by ligand internalization, fluorescently labelled ligands can be used to track ligand internalization and thus receptor activation. A secretin variant containing the near-infrared fluorescent dye IDCC was synthesized and used to stimulate SCTR in β-arr2-GFP/SCTR-coexpressing U2OS cells. As Fig. 3C demonstrates, the subcellular distribution patterns of β-arr2-GFP (green) and of secretin-IDCC (red) after ligand addition coincide to a large degree, yielding a high proportion of yellow vesicle structures seen in the overlay of both fluorescence channels. The functional equivalence of secretin and secretin-IDCC is shown in Fig. 3D. The EC_50_ values determined by vesicle quantitation in the green channel (β-arr2-GFP) and in the red channel (secretin-IDCC) proved to be very close: 3.02 ± 0.12 nM and 1.45 ± 0.54 nM, respectively. Consequently, the reporter tag IDCC did not interfere with secretin function in this assay. Similarly, secretin-IDCC was competitively displaced by secretin (Fig. 3D, inset), allowing for the assay to be run in antagonist mode.

### 3.3. Alanine walk

To gain insight into the contribution of amino acids to the agonistic activity of the peptide, every residue in human secretin was individually substituted by Ala. All 27 peptide variants including the wild-type with a native Ala^17^ were tested in the β-arr2-GFP translocation assay to measure potency and intrinsic activity (Figure 4 first row, Supplementary Table 1). Amino acid residues with primary importance for agonistic activity were almost exclusively found in the N-terminal region of secretin (Fig. 4A). The strongest effect was seen with the Asp^3^ substitution, resulting in a 500-fold increase in the observed EC_50_. Exchange of His^1^, Phe^6^, Thr^7^ and Leu^10^ increased the EC_50_ value by a factor of 60-70. All other residues were exchanged by Ala without a huge reduction in potency. On the other hand, effects on intrinsic activity were less pronounced. Intrinsic activity dropped to 60 % with the substitution of Asp^3^ and Phe^6^, while others were similar to the wild type (Fig. 4B, first row).

**Figure 4:**
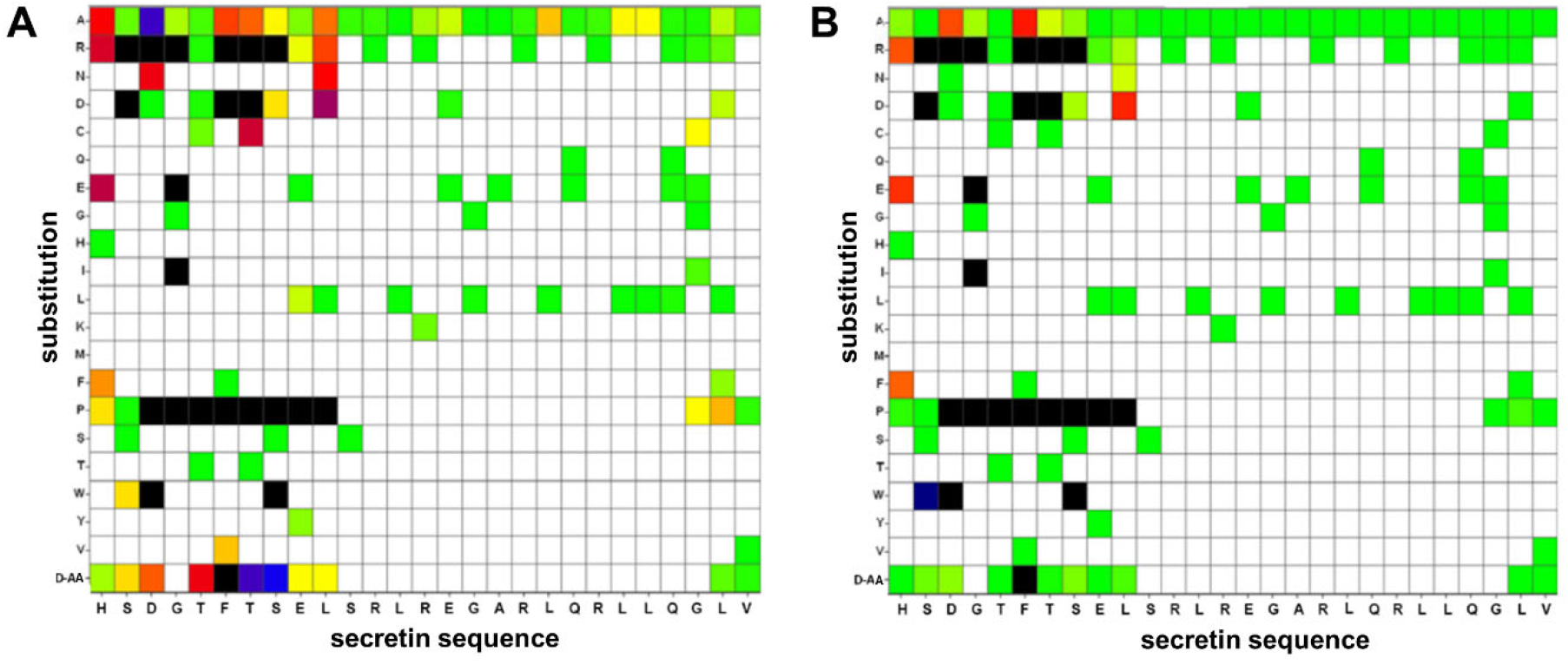
Effects of single amino acid substitutions in secretin on potency (A) and intrinsic activity (B). Single amino acids of human secretin were substituted by other natural or the respective D-amino acids activity measured by β-arr2-GFP translocation. The x-axis displays the native sequence of secretin while the y-axis shows the individual amino acid exchange. D-AA indicates a substitution by the respective amino acid in D-conformation. **A)** Amino acid substitutions depicted in green showed no elevation in the resulting EC_50_ value (EC_50_ of native secretin: 3.26 ± 0.80 nM), yellow: EC_50_ up to 20 nM, red: EC_50_ up to 200 nM, blue: EC_50_ up to 2000 nM, black: not detectable. White areas indicate variants not studied. **B)** Amino acid substitutions labeled in green display an intrinsic activity comparable to unmodified human secretin (100%). The course from yellow over red, blue towards black indicates deterioration of the intrinsic activity: yellow: efficacy below 80%, red: below 60%, blue: below 40%, black: not detectable. White areas indicate variants not studied. All values shown represent the statistical mean of at least three independent experiments performed in duplicates.

### 3.4 Additional single substitution variants

To expand the basic insights gained by the alanine walk, a further 69 single amino acid substitution variants were evaluated using the same assay. Residues for exchange were chosen to represent different physicochemical properties of the side chains as well as D-amino acids, with most of the substitutions focusing on the N-terminal portion of the peptide. (Fig. 4, Supplementary Table 1 and 2). The most prominent reductions in EC_50_ and intrinsic activity were caused by the introduction of proline and arginine. The aminoterminal His^1^, e.g., was substituted by D-His without noticeable reduction of potency, while Pro decreased the EC_50_ by a factor of 10. This adverse effect was further enhanced by the introduction of the charged amino acids Glu and Arg, displaying a 200-fold EC_50_ reduction (Fig. 4A). These modifications also lead to a marked reduction of the intrinsic activity (Fig. 4B). An exchange of Gly^2^ to Pro^2^ did not diminish the peptide’s activity, while substitution with Arg, Asp and Trp had severe negative effects on both potency and intrinsic activity. Asp^3^ proved to be highly susceptible to substitution. D-Asp and Asn elevated the EC_50_ by a factor of 50- to 100-fold while exchanges for Arg, Pro and Trp completely abolished agonistic activity. Position 4 likewise only tolerated Ala as substitute in order for the peptide to maintain functional. Threonine on position 5, in contrast, showed little vulnerability with the exception of Pro and D-Tyr.

The phenylalanine in position 6 was known to be highly conserved in peptide ligands of all class B GPCRs [30]. Indeed, all tested substitutions deteriorated EC_50_ values, most of them also intrinsic activity. Val at this position led to a drastic reduction in the EC_50_, whereas all other substitutions reduced potency below the detection limit. Threonine on position 7 seemed to play a crucial role for an agonistic reaction, too. Any of the tested substitutions at this residue led to a pronounced loss of potency, with Arg, Asp and Pro compromising intrinsic activity. The serine at position 8 was replaced by Asp with a slight elevation of the EC_50_ while Arg, Pro, Trp and D-Ser were not favorable. Apart from substitution with Pro, glutamic acid on position 9 was proven rather insensitive to the tested exchanges. Leu^10^ showed moderate to strong negative effects as a result to substitution. Most positions examined displayed EC_50_ changes in a range of 45-300-fold. Beyond position 11, secretin proved to be highly tolerant towards the examined single amino acid substitutions.

### 3.5. Truncation variants

To further elucidate the contribution of amino- and carboxyterminal domains of secretin to the peptide’s biological activity, the wild-type peptide was successively truncated from either end and variants were evaluated for potency and intrinsic activity. Using the β-arr2-GFP translocation assay, only a small number of activity profiles could be derived within a concentration range up to 10 μM, as truncated peptides dramatically lost activity (data not shown). In direct comparison, cAMP production measurements proved to be of higher sensitivity, displaying EC_50_ values several orders of magnitudes lower than those derived from β-arr2-GFP translocation experiments. For secretin e.g., the β-arr2-GFP EC_50_ was found to be 3.26 ± 0.8 nM, while that for cAMP production was 0.0012 ± 0.0003 nM. Therefore, characterization of the truncated peptide variants was carried out by intracellular cAMP measurements. Figure 5 displays normalized EC_50_ values of truncated peptide variants in direct comparison to native human secretin. Progressive truncation on either Cor N-terminus increasingly lowered the potency (Figure 5, Supplementary Table 2). As expected, small aminoterminal truncations yielded a stronger effect than similar changes at the C-terminus.

**Figure 5:**
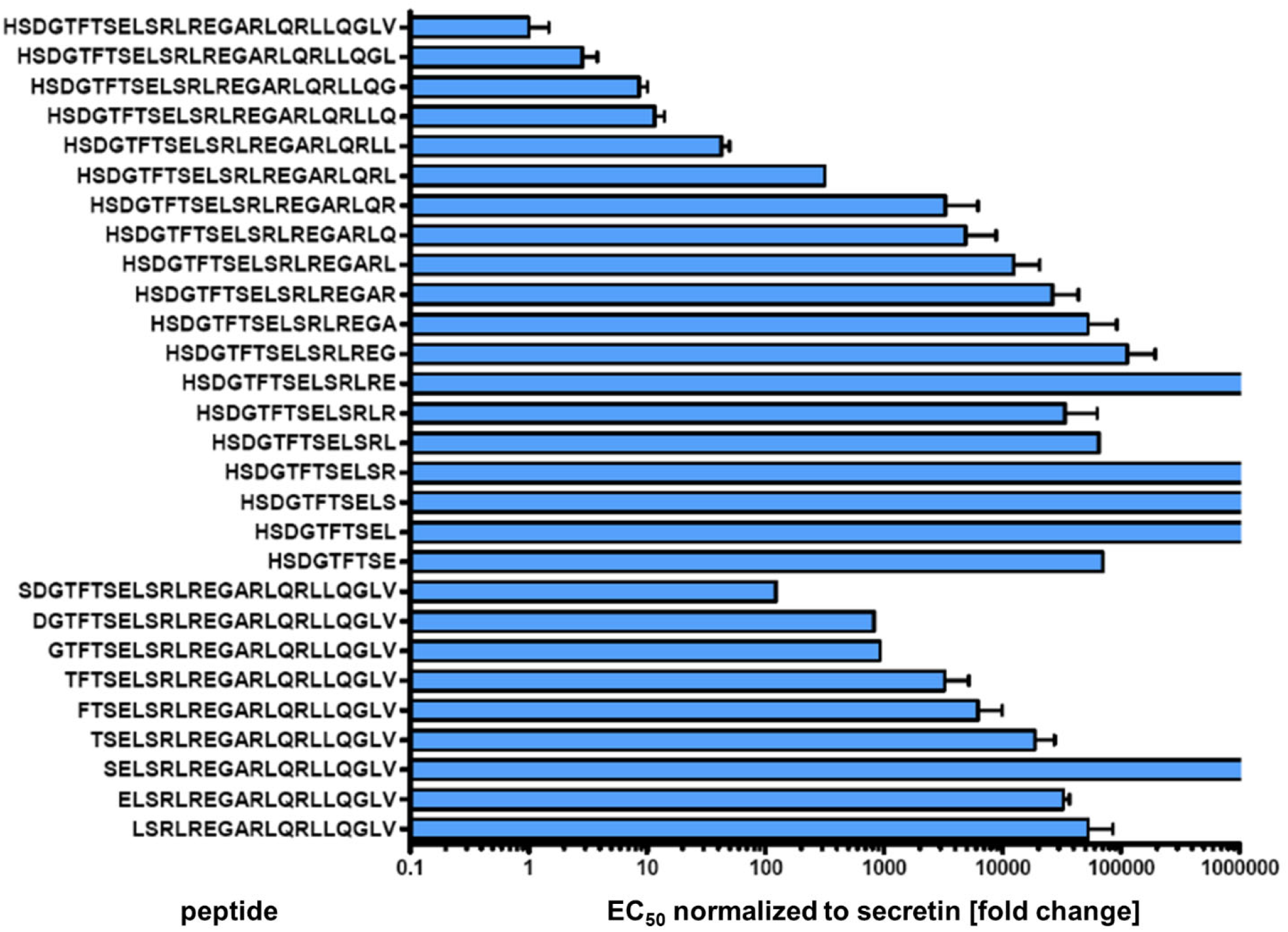
Intracellular cAMP activation of SCTR by N- or C-terminal truncated secretin variants Individual peptide sequences are given at the y axis. EC_50_ values were normalized to wild-type secretin (upper row). Values result from at least three independent experiments performed in duplicates.

### 3.6. Characterization of antagonistic properties of secretin analogs

To investigate whether these N-terminal truncation variants (peptides 115-123) with drastic EC_50_ reduction would show antagonistic properties, they were tested in the β-arr2-GFP translocation assay run in antagonist mode. Indeed, most of these aminoterminal truncation variants demonstrated antagonism with IC_50_ values of approximately 2 μM (Table 2).

**Table 2.**
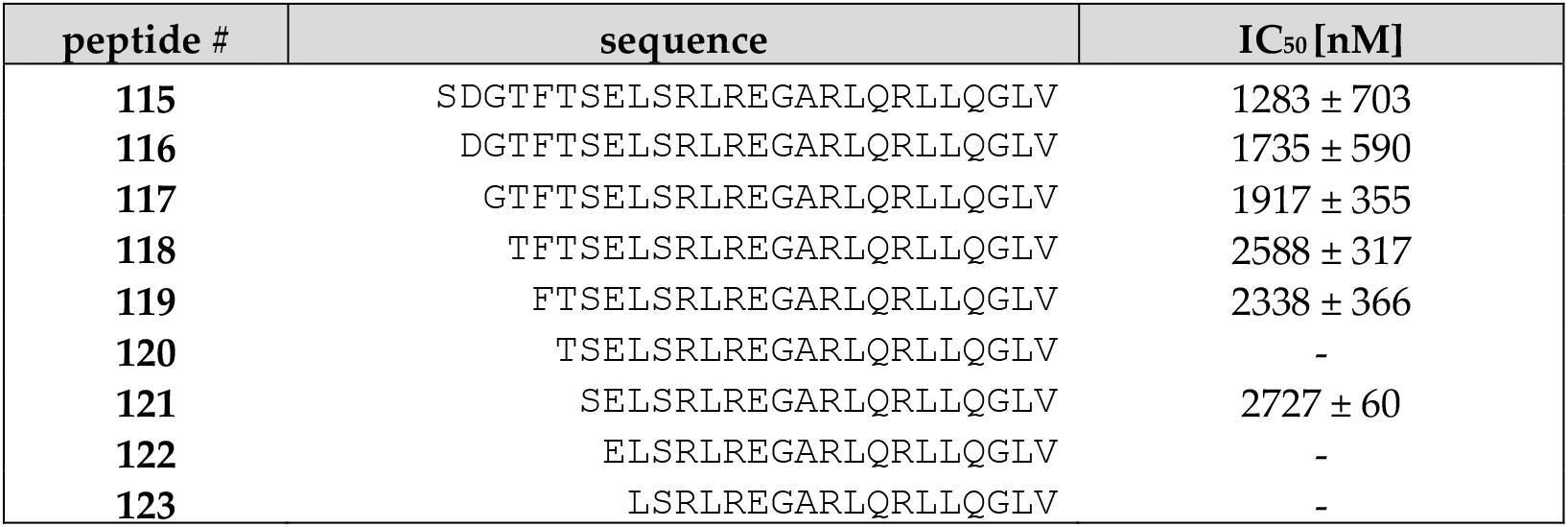
SCTR antagonism of secretin variants truncated at the N-terminus, assayed via inhibition of β-arr2-GFP translocation. (-): no detectable reaction. All displayed IC_50_ values result from at least 3 independent experiments performed in duplicates.

Of those variants, peptide 119 was one of the shortest with measurable antagonistic properties (Table 2). In an attempt to enhance its antagonistic activity, the N-terminus was elongated by one charged amino acid. Indeed, peptides 124 and 125, extended by Arg or D-Arg at the N-terminus, showed a twofold improvement in the IC_50_ to about 1 μM (Table 3). In order to probe the D-Arg-extended variant for further enhancement in antagonism, another alanine walk based on this peptide was carried out. While six variants lost their antagonistic capacity and another seven showed a higher IC_50_ than peptide 125, seven of the peptides had retained or improved their antagonistic capacity. Among them, peptide 136 was found to have an IC_50_ of 309 ± 74 nM.

**Table 3:**
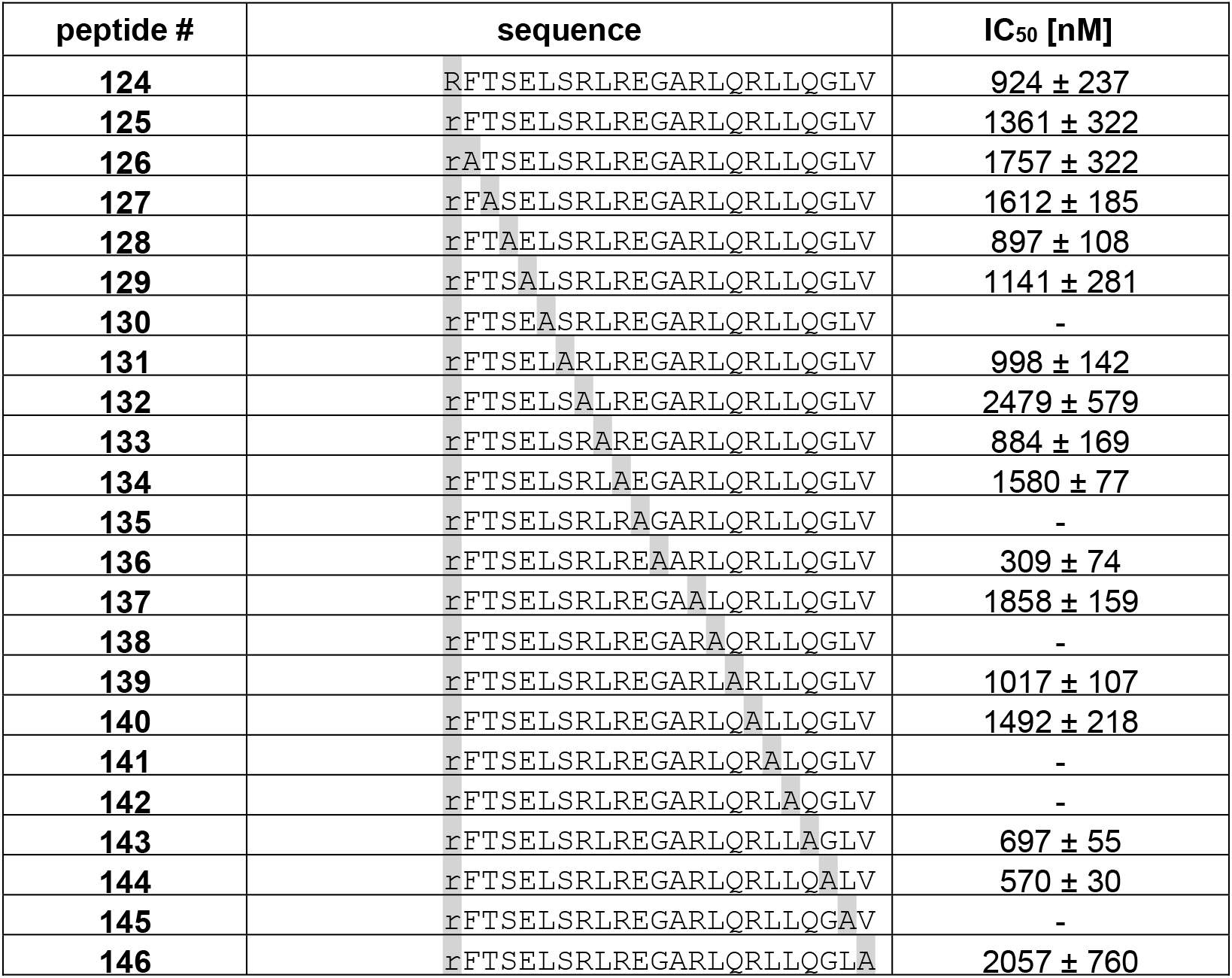
Impact of single amino acid exchanges in N-terminal truncation variants of secretin, as assayed by inhibition of β-arr2-GFP translocation. Substituted amino acids are highlighted in grey. (-): no measurable reaction, r: D-Arg. All displayed values are the result of at least n=3 independent experiments performed in duplicates.

### 3.7 Further characterization of selected secretin variants

For five selected peptides, the characterization was further extended by determining radioligand binding and cAMP response data. Three of the Ala-modified N-terminal truncations (peptides 136, 143 and 144) as well as peptide 120 and wild-type secretin (peptide 1) were included. Peptide 120, a variant with a deletion of the first six amino acids, secretin(7-27), did not show any detectable antagonism in the β-arr2-GFP translocation assay (Table 4). On the other hand, peptides 136, 143 and 144 demonstrated an IC_50_ of 309 ± 74 nM, 697 ± 55 nM, and 570 ± 30 nM, respectively (Figure 6A, Table 4). None of these three peptides showed any detectable activation (agonistic properties) in the β-arr2-GFP translocation assay (Figure 6B). Surprisingly, these peptides showed different properties regarding cAMP production. Here, no antagonistic activity was observed (data not shown). In contrast, the three peptides proved to be partial agonists in the cAMP assay, with EC_50_ values around 30 nM and intrinsic activities around 30% (Table 4, Fig. 6C). In comparison, wild-type secretin showed an EC_50_ of 0.1 ± 0.03 pM in this assay. Interestingly, the unmodified truncated secretin(7-27) retained a high intrinsic activity (90.6 ± 11.5 %) but only residual potency in the cAMP response (3366 ± 1570 nM)

**Figure 6:**
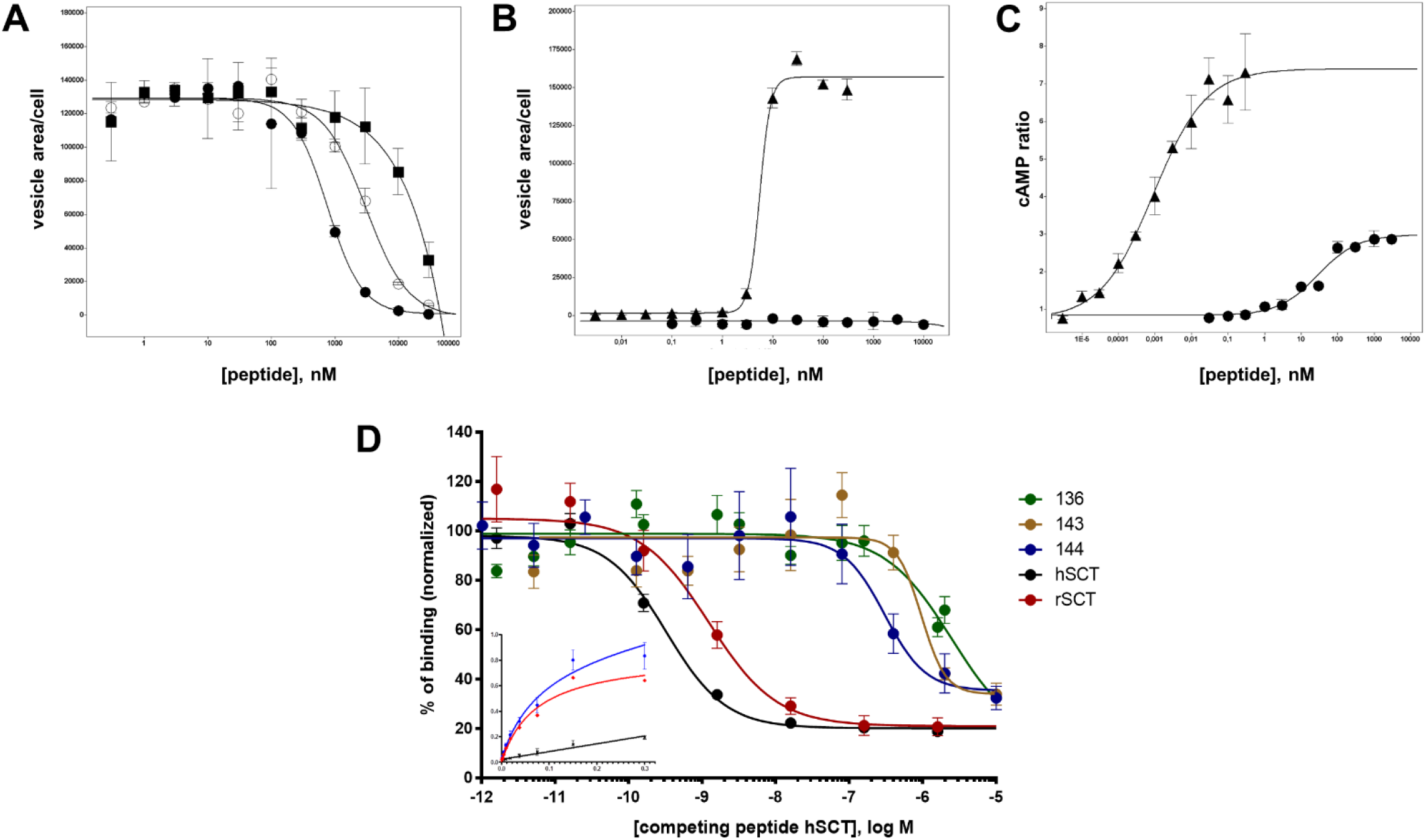
Antagonist and agonist assays of wild-type secretin and selected secretin variants. **A)** Peptides 144, 117, and 122 were analyzed for SCTR antagonism in the β-arr2-GFP translocation assay: peptide 144, closed circles (●); peptide 117, empty circles (∎); peptide 122, closed squares (∎). **B)** SCTR agonism measured in the β-arr2-GFP translocation assay: wild-type secretin, closed triangles (▲); peptide 144, closed circles (●). **C)** SCTR agonism measured in the cAMP GloSensor assay: wild-type secretin, closed triangles (▲); peptide 144, closed circles (●). **D**) Competition binding of radioligand and secretin variants on U2OS-SCTR membrane isolates. I^125^-secretin was incubated with purified membranes of SCTR-overexpressing cells. IC_50_ for human secretin was calculated at 0.325 nM (black), 1.231 nM for rat secretin (red), 2472 nM for peptide 136 (green), 963 nM for peptide 143 (yellow) and 304 nM for peptide 144 (blue). Displayed binding data represent at least three independent experiments, performed in triplicates. Inset: saturation binding curves obtained with I^125^-secretin used in the aforementioned competition experiments: blue curve: total binding, black curve: unspecific binding, red curve: specific binding (K_D_ = 74 pM).

**Table 4:**
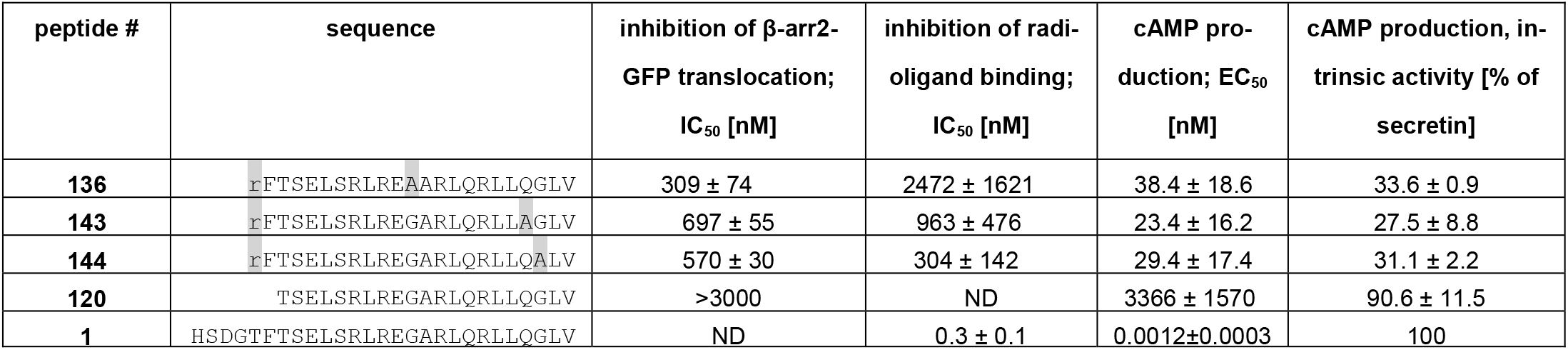
SCTR antagonism measured as inhibition of β-arr2-GFP translocation, competition radioligand binding and SCTR agonism measured by the GloSensor cAMP assay performed on selected secretin variants. Substituted amino acids in comparison to human secretin (peptide 1) are highlighted in grey. All displayed values are derived from at least 3 independent experiments performed in duplicates. ND: not determined.

### 3.8 Analysis of binding affinity

To compare these results with radioligand binding data, a modified human secretin was labelled with ^125^iodine and HPLC-purified radiopeptide was used in a binding assay on isolated membranes of an SCTR-overexpressing cell line, followed by scintillation counting. Mean scintillation values displayed half-maximal displacement at competitor concentrations of 0.325 nM for unmodified human secretin and 1.231 nM for the structurally similar rat secretin (Fig. 6D). Vasoactive intestinal polypeptide, which has been known to partially interact with SCTR by its amino terminal end, displayed an IC_50_ approximately 2300-fold lower than the native ligand. Affinities for peptides 143 and 144 (but not for 136) were in the same nominal range as determined for the β-arr2-GFP translocation assay, namely 2472 ± 1621 nM, 963 ± 476 nM, and 304 ± 142 nM (Fig. 6D, Table 4).

## 4 Discussion

Personalized medicine and precision oncology represent concepts that address medical needs resulting from limited efficacy and specificity of current treatment options [47,48]. Both concepts aim at improving patient survival and quality of life by considering interindividual variation in genetic aberrations, protein expression, metabolic regulation, immune activity and other disease drivers [49,50]. One of the strategies pursued to personalize disease management in oncological settings is called tumor targeting: utilizing the overexpression of cell surface receptors in tumors for the accumulation of high-affinity ligands equipped with suitable effectors, typically radionuclides, for imaging or therapy [51,52]. Both pancreatic ductal adenocarcinoma as well as esophageal cancer represent tumor entities with inadequate treatment options and unfavorable patient prognosis. Five-year survival rates are 20% for esophageal cancer, and 11% for pancreatic cancer [53]. Thus, there is a high medical need to improve therapeutic options for these tumors, by e.g., taking advantage of cell surface receptor overexpression and the use of optimized tracer molecules.

In this study, the expression of secretin receptor (SCTR) was characterized by immunohistochemistry of paraffin-embedded human tissue sections. To the best of our knowledge, this is the first description of an antibody to detect SCTR expression in human tumor tissues. Many G protein-coupled receptors suffer from a lack of highly specific antibodies [54–57]. This has only recently been addressed by the development of improved commercially available immunoglobulin preparations. By employing suitable positive and negative controls, the antibody applied here was extensively characterized prior to use on target tissues (Supplementary Figure 1). After successful validation, the SCTR antibody was utilized to detect SCTR expression in normal as well as tumor tissues. For all three subtypes of esophageal cancer, a strong SCTR overexpression was detected, most prominent in ESCC. Intriguingly, the plasma membrane localization of SCTR was found to be conserved in overexpressing tumors, providing for an opportunity to target these cell-surface receptors with appropriate tracers. This retention of plasma-membrane localization was also found in PDAC tissues, where high SCTR overexpression was found in 65 of 70 investigated PDAC tissues, while islet cell tumors and chronic pancreatitis proved to be low in SCTR expression. SCTR expression in PDAC has previously been reported, yet not as an overexpression: SCTR was detected in only 5 out of 12 samples, with tumor-free pancreas always positive [58], or as markedly reduced in tumors as compared with normal pancreas [39]. Both these studies employed autoradiography with ^125^I-secretin, while this work relied on an antibody and immunohistochemistry for the detection of SCTR. Whether the differences observed result from this methodological difference remains to be determined, as similar results were obtained with both techniques e.g., for normal pancreas and liver [39,40].

Interestingly, high SCTR overexpression was demonstrated in tumor tissues of all histopathological gradings, with only slightly lower values for less differentiated tumors. This contrasts with the literature on GPCR overexpression in other tumors such as neuroendocrine neoplasms, where overexpression of somatostatin receptors and the receptor for gastric inhibitory polypeptide may be variable or reduced in G3 tumors in contrast to G1 and G2 tumors [59,60]. In colorectal carcinoma (CRC), a massive methylation of the SCTR gene with lower methylation levels in precancerous lesions (polyp, adenoma) and highest levels in carcinoma has recently been reported [61]. Consequently, and in contrast to the findings in this study for PDAC and esophageal cancer, SCTR expression in CRC may be strongly reduced rather than upregulated. In three PDAC cell lines, the SCTR promotor had been found unmethylated, stressing different gene expression regulation between CRC and PDAC [62]. Apart from the requirement of overexpression in tumor tissue, the utility of a cell surface receptor for targeting depends on its preferably low expression in other organs, in particular those relevant for contrast in imaging modalities such as single-photon emission tomography (SPECT) and positron emission tomography (PET). In addition, organs typically involved in radiopharmaceutical toxicity such as kidney and bone marrow should be low in the expression of the target receptor. The expression of SCTR in certain cell populations within the kidney as well as the wide SCTR expression in ductal and other epithelia should thus be further evaluated for their potential contribution to background imaging signals as well as for potential dose-limiting toxicity in PRRT.

Due to their vulnerability towards proteolytic degradation, most peptide hormones have short half-lives of only a few minutes. For secretin, a half-life of 2.5-4 minutes has been reported [63–66]. To chemically stabilize a peptide ligand by the introduction of artificial amino acids, the amino acid positions crucial for its activity need to be identified. The experimental strategy included design and synthesis of a library of secretin variants that would contain single amino acid substitutions as well as truncations on either end. To characterize a large number of individual peptides in a quantitative concentration-response setup, a β-arr2-GFP translocation assay was established in U2OS cells coexpressing human SCTR and β-arr2 coupled to GFP. As has been shown before, the intracellular translocation of GFP-labeled β-arrestin upon ligand application can be utilized as a sensitive, robust, and universal readout system for screening G-protein-coupled receptors [46]. Likewise, for rat SCTR overexpressed in HEK293 cells, ligand-dependent translocation of over-expressed arrestin-GFP had been demonstrated [34]. Using an approach utilizing automated microscopy and image analysis, 146 secretin variants were analyzed for their agonistic (EC_50_, intrinsic activity) and/or antagonistic (IC_50_) capacity. As an additional tool to monitor receptor activation, internalization of a fluorescently labeled secretin was also followed in the microscopic system. Both β-arrestin translocation and ligand internalization showed very similar quantitative results.

This study utilized a library approach to probe the functional requirements on within human secretin’s primary structure on cells expressing human SCTR. The first step in this approach was the analysis of Ala substitutions of all amino acid position except for the natural Ala^17^. A related investigation has previously been published for rat secretin and rat SCTR [20]. Results were similar in that both studies found amino acid positions 1,7, 8, 10, 19, 22, and 23 to be most critical for biological activity. However, they also differed in a few other positions: while this study found Asp^3^ to be most susceptible to Ala substitution and identified Phe^6^ as critical in human secretin, Dong et al found Gly^4^ and Glu^9^ to be of crucial importance for rat secretin to activate rat SCTR. While the authors did not report the Ala substitutions’ capacity to modulate intrinsic activity, the current study revealed Ala substitution of Asp^3^ and Phe^6^ to be most detrimental to the peptide’s intrinsic activity. Similarly, substitution of positions 1, 3, 4, 7, 8, 15, and 19 of rat secretin by a phenolic residue had been shown previously to impair activation of rat SCTR [67], as was the introduction of Phe^1^, Trp^3^ and Trp^8^ for human SCTR in the results presented here. These results are also in agreement with the recently published cryo-EM structure of a secretin-SCTR-Gs complex [21] that demonstrated close interaction of secretin amino acids 1, 3, 8 and 15 with the receptor, while no proximity was identified for the other activity-related positions mentioned above. Further substitution variants focusing on the N-terminal part of secretin generally confirmed the high structural requirements for SCTR activation, with the notable exception that the exchange of Gly^2^ to Pro^2^ did not diminish the peptide’s activity. Phe^6^, known to be highly conserved in peptide ligands of all class B GPCRs [30], proved to be most sensitive to exchange, even for Tyr.

Early studies of secretin had shown that its biological activity resides particularly in the aminoterminus. A number of carboxyterminal fragments, among them secretin(7-27), had been shown to act as weak SCTR antagonists [24,68]. Secretin(2-27) displayed a severe reduction on cAMP synthesis on isolated pancreatic membranes, while the truncation to amino acids(7-27) completely abolished agonistic effects of the peptide variant [24]. Despite their lack of activation, these truncated peptides were able to displace native secretin at high concentrations [24,25]. Secretin(7-27) did not show any detectable antagonism in the β-arr2-GFP translocation assay in this study, however. Using guinea pig pancreatic acini, a number of reduced peptide bond pseudopeptide antagonistic analogues of porcine secretin [25] with improved specificity for SCTR have previously been developed. The most active compound showed an IC_50_ of 2700 ± 500 nM in the cAMP inhibition assay, with no measurable agonism. In the radioligand competition binding assay, this compound had a Ki of 4400 ± 400 nM. Starting from the truncated sequence of rat secretin(5-27), substituted variants have been generated in a previous study [16]. The variant with the highest affinity of 23 nM was [I17, R25]-secretin(5-27). In the current study, the most potent antagonist showed an IC_50_ of 309 ± 74 nM in the β-arr2-GFP translocation assay on human SCTR at a moderate affinity of 2472 ± 1621 nM and a second one with an antagonistic activity (IC_50_) of 570 ± 30 nM and an affinity of 304 ± 142 nM. Further investigating the contribution of sequence motifs for the activation of individual signaling pathways (cAMP production, calcium release, β-arrestin activity) may help identify biased SCTR ligands [69]. Recently, the rational development of a SCTR antagonist with multiple substitutions showing an affinity of 4 nM has been reported [26]. In view of the emerging opportunities to utilize SCTR ligands for molecular imaging and targeted therapy in gastrointestinal cancer, our knowledge about structural requirements in secretin for both binding and activation needs to be expanded. This study has further delineated the structure-activity relationship in this peptide ligand and reported a number of new lead structures for β-arrestin antagonism in SCTR. Future studies will have to demonstrate the utility of secretin analogues as tracers for molecular imaging and targeted therapy of cancer.

## Supplementary Materials

The following supporting information can be downloaded at: xxx Figure S1: SCTR antibody validation; Figure S2: SCTR IHC score for different histological gradings; Figure S3: SCTR staining in normal tissues; Table S1: Pharmacological properties of wild-type secretin and secretin variants, part 1; Table S2: Pharmacological properties of secretin variants, part 2.

## Author Contributions

Conceptualization, C.G., B.W. and A.K.; methodology, A.K., S.A., and L.N.; formal analysis, A.K., S.A., and L.N.; investigation, A.K., S.A., and L.N.; writing—original draft preparation, C.G..; writing-review and editing, C.G. and A.K..; visualization, A.K. and C.G..; supervision, C.G. and B.W.; funding acquisition, C.G. All authors have read and agreed to the published version of the manuscript.

## Funding

This research was funded by Bundesministerium für Bildung und Forschung (BMBF), grant numbers IP614 and IPT614A to CG.

## Institutional Review Board Statement

Not applicable.

## Data Availability Statement

Numerical, machine-readable data for this study have been deposited in an open data repository for public access: https://doi.org/10.5281/zenodo.5925746

## Acknowledgments

The authors wish to thank Ines Eichhorn and Yvonne Giesecke for excellent technical support. The authors acknowledge support by the Molecular Cancer Research Center (MKFZ), Charité - Universitätsmedizin Berlin, 13353 Berlin, Germany

## Conflicts of Interest

The authors declare no conflict of interest.

## Notes

https://doi.org/10.5281/zenodo.5925746

### Competing Interest Statement

The authors have declared no competing interest.

