## Supplement for "Secretin receptor as a target in gastrointestinal cancer: expression analysis and ligand development"

*Antibody validation*

A commercially available SCTR antibody (HPA007269, Sigma-Aldrich) was validated for specificity and subcellular staining. An epitope tag antibody (TAG) detected expression of SCTR equipped with this epitope tag and of the related glucagon-like receptor 2 (GLP-2R), respectively, in lysates in a Western blot. The SCTR antibody, however, detected only the expressed SCTR-TAG protein, proving specificity of the antibody for SCTR (Supplementary figure 1A). Likewise, U2OS cells stably expressing either SCTR or GLP-2R were stained with the SCTR antibody in immunofluorescence. Applying both permeabilizing and non-permeabilizing conditions, the SCTR antibody only yielded a signal in SCTR-expressing, not GLP-2R-expressing cells, confirming its high specificity. In addition, the localization of the SCTR staining at the plasma membrane was found to be consistent with the detection of a heptahelical receptor like SCTR (Supplementary figure 1B).

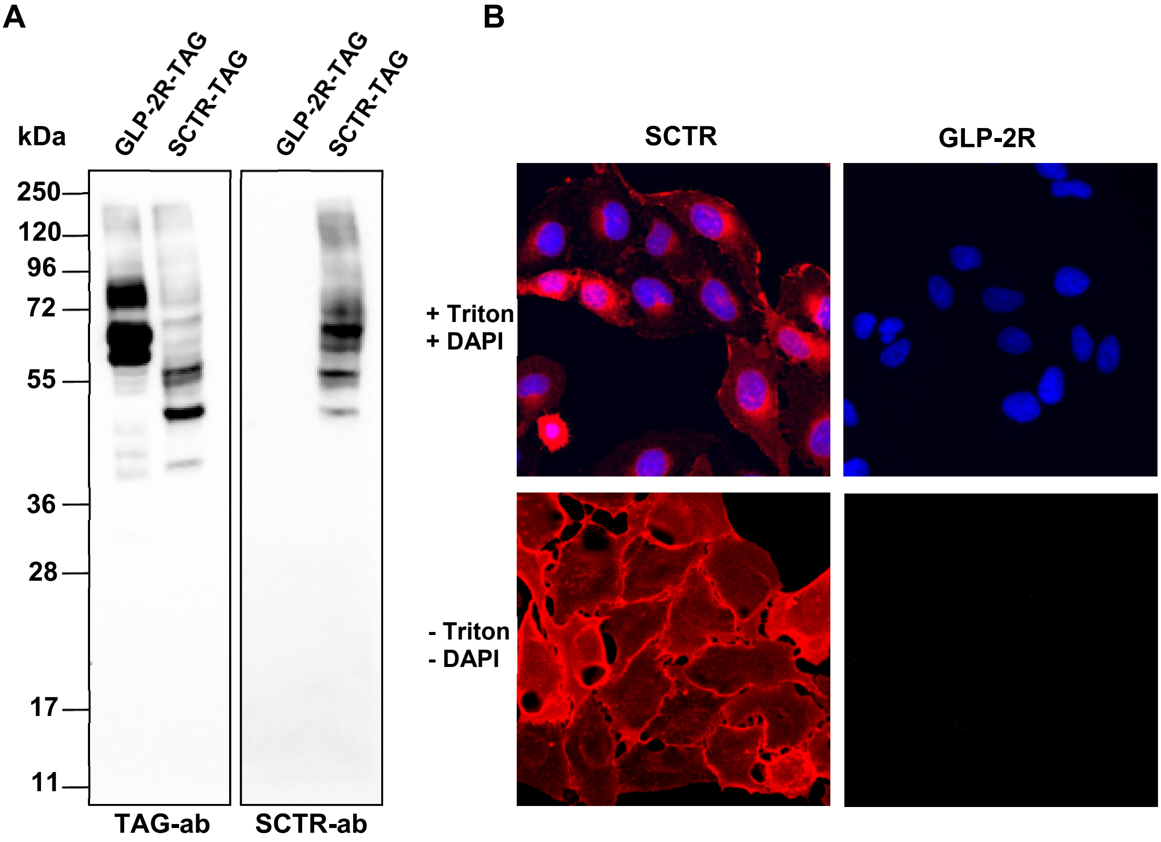


**Supplementary figure 1:** Validation of the antibody against SCTR. (**A**) Western blot: U2OS cells were transiently transfected with a plasmid encoding GLP-2R equipped with an epitope tag (GLP-2R-TAG) or with a plasmid encoding SCTR with the same tag (SCTR-TAG), and lysed 24 h later. Lysates were separated by SDS-PAGE and blotted onto a nitrocellulose membrane. The left part of the membrane was incubated with an antibody against the epitope tag (TAG-ab), the right part of the membrane with the antibody against SCTR (SCTR-ab, 0.16 µg/ml, Sigma-Aldrich). Detection was performed using a POD-coupled goat anti-mouse or goat anti-rabbit antibody and SuperSignal West Dura development substrate (PIERCE). (**B**) Immunofluorescence: U2OS cells stably expressing SCTR and GLP-2R, respectively, were fixed and partially permeabilized with 0.1% Triton-X100. Cells were stained with the antibody against SCTR (0.4 µg/ml, Sigma-Aldrich) and a Cy3-labeled goat anti-rabbit antibody. The nuclei of the permeabilized cells were additionally stained with DAPI. Images were acquired on the InCell Analyzer 1000 (GE Healthcare).

*SCTR IHC score for different histological gradings*

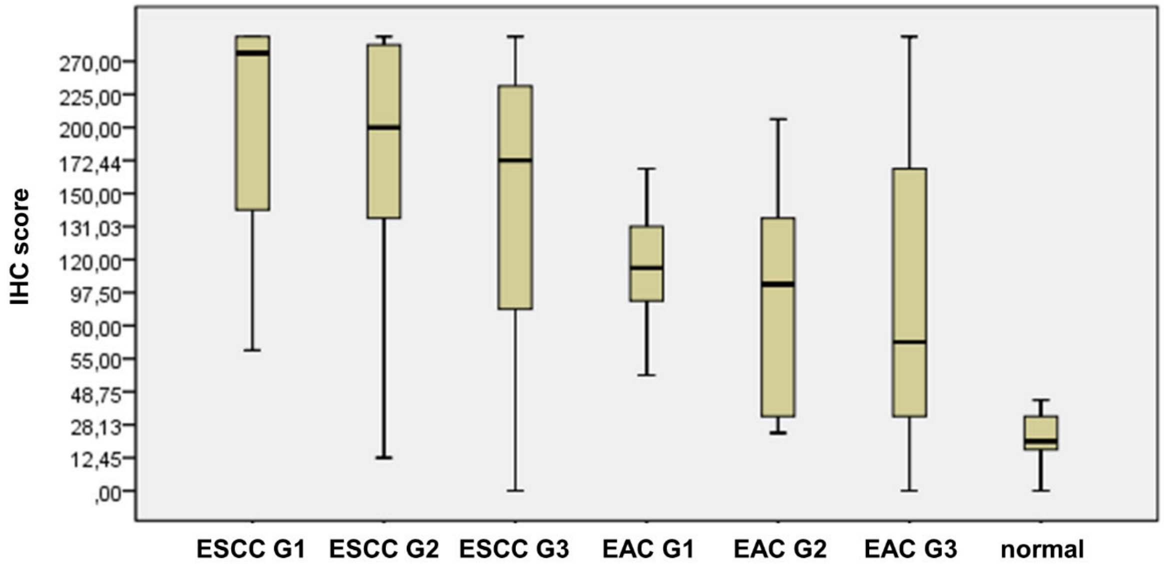


**Supplementary figure 2:** Boxplot of the IHC score of SCTR antibody staining in normal and cancerous esophageal samples. ESCC and EAC IHC scores were plotted according to the histopathological grading (G1, G2 or G3) of the tumors. For comparison, the IHC score for normal esophageal tissue is shown.

*SCTR staining in normal tissues*

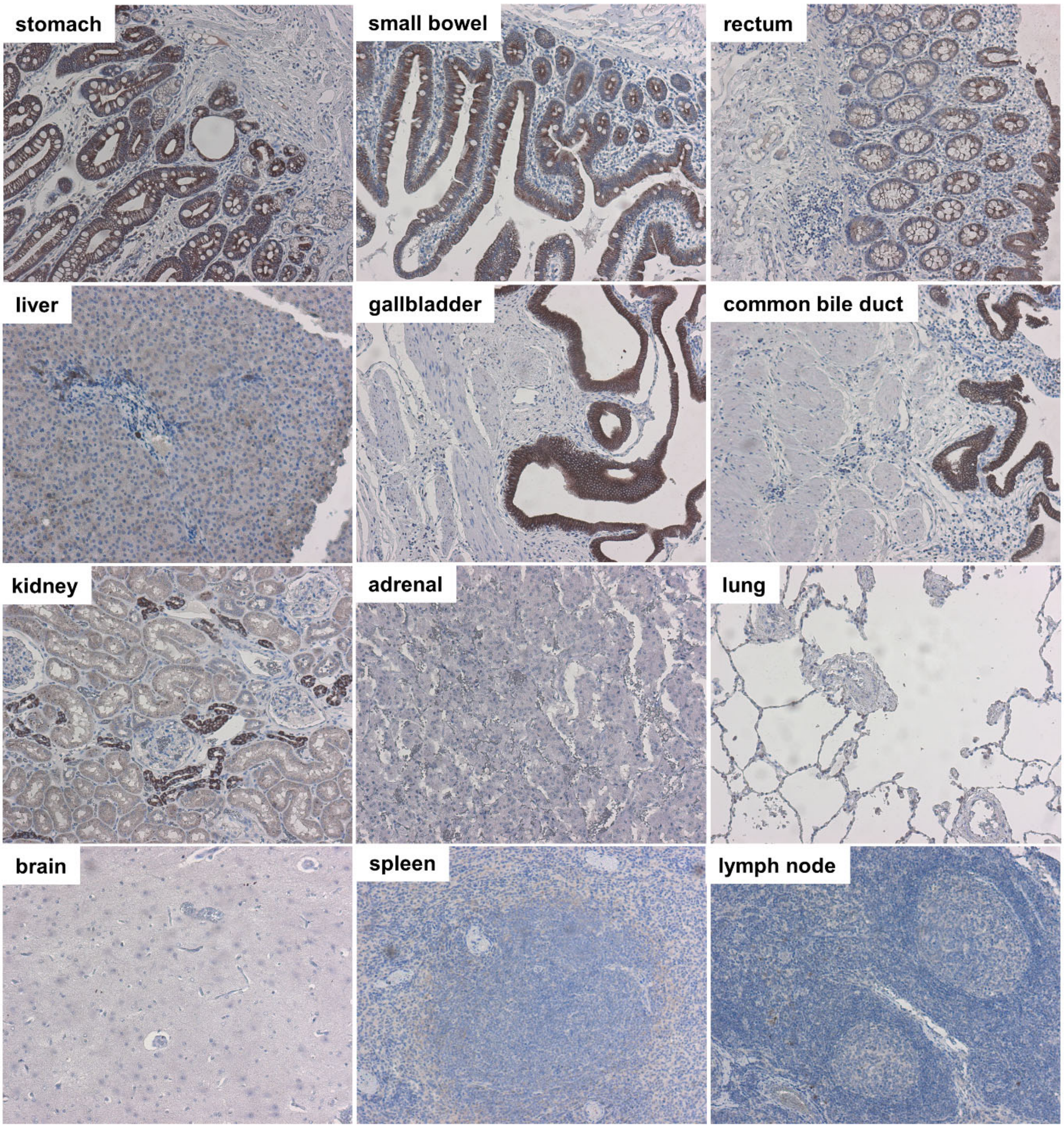


**Supplementary figure 3: SCTR immunohistochemistry of normal tissues.** As for esophageal and pancreatic normal and pathological samples, a number of normal tissues were stained for SCTR, too. These included samples from stomach, small bowel, rectum, liver, gallbladder, common bile duct, kidney, adrenal, lung, brain, spleen and lymph node, each n=1. Microscopic images were taken using a 10x objective.

### Supplementary table 1

Pharmacological properties of wild-type secretin and secretin variants, part 1.

| peptide | sequence 1 | 2 | 3 | 4 | 5 | 6 | 7 | 8 | 9 | 10 | 11 | 12 | 13 | 14 | 15 | 16 | 17 | 18 | 19 | 20 | 21 | 22 | 23 | 24 | 25 | 26 | 27 | β-arrestin2-GFP agonism potency [nM] | SD [nM] | efficacy [%Sec] | SD [%Sec] | cAMP agonism potency [nM] | SD [nM] |
|---|---|---|---|---|---|---|---|---|---|---|---|---|---|---|---|---|---|---|---|---|---|---|---|---|---|---|---|---|---|---|---|---|---|
| 1 | H | S | D | G | T | F | T | S | E | L | S | R | L | R | E | G | A | R | L | Q | R | L | L | Q | G | L | V | 3,26 | 0,80 | 100,00 | | | |
| 2 | A | S | D | G | T | F | T | S | E | L | S | R | L | R | E | G | A | R | L | Q | R | L | L | Q | G | L | V | 233,86 | 119,49 | 89,10 | 4,71 | | |
| 3 | H | A | D | G | T | F | T | S | E | L | S | R | L | R | E | G | A | R | L | Q | R | L | L | Q | G | L | V | 9,21 | 6,84 | 102,60 | 4,78 | | |
| 4 | H | S | A | G | T | F | T | S | E | L | S | R | L | R | E | G | A | R | L | Q | R | L | L | Q | G | L | V | 1.577,92 | 1.386,13 | 65,88 | 24,59 | | |
| 5 | H | S | D | A | T | F | T | S | E | L | S | R | L | R | E | G | A | R | L | Q | R | L | L | Q | G | L | V | 14,01 | 7,37 | 87,17 | 4,35 | | |
| 6 | H | S | D | G | A | F | T | S | E | L | S | R | L | R | E | G | A | R | L | Q | R | L | L | Q | G | L | V | 6,99 | 4,31 | 104,45 | 8,49 | | |
| 7 | H | S | D | G | T | A | T | S | E | L | S | R | L | R | E | G | A | R | L | Q | R | L | L | Q | G | L | V | 155,16 | 30,79 | 61,83 | 4,92 | | |
| 8 | H | S | D | G | T | F | A | S | E | L | S | R | L | R | E | G | A | R | L | Q | R | L | L | Q | G | L | V | 131,37 | 48,42 | 83,88 | 12,51 | | |
| 9 | H | S | D | G | T | F | T | A | E | L | S | R | L | R | E | G | A | R | L | Q | R | L | L | Q | G | L | V | 24,29 | 12,76 | 88,08 | 10,83 | | |
| 10 | H | S | D | G | T | F | T | S | A | L | S | R | L | R | E | G | A | R | L | Q | R | L | L | Q | G | L | V | 10,87 | 3,36 | 100,17 | 11,42 | | |
| 11 | H | S | D | G | T | F | T | S | E | A | S | R | L | R | E | G | A | R | L | Q | R | L | L | Q | G | L | V | 120,52 | 23,18 | 96,50 | 9,58 | | |
| 12 | H | S | D | G | T | F | T | S | E | L | A | R | L | R | E | G | A | R | L | Q | R | L | L | Q | G | L | V | 5,76 | 1,89 | 101,60 | 10,47 | | |
| 13 | H | S | D | G | T | F | T | S | E | L | S | A | L | R | E | G | A | R | L | Q | R | L | L | Q | G | L | V | 6,06 | 2,04 | 100,37 | 18,43 | | |
| 14 | H | S | D | G | T | F | T | S | E | L | S | R | A | R | E | G | A | R | L | Q | R | L | L | Q | G | L | V | 2,71 | 1,46 | 99,50 | 12,76 | | |
| 15 | H | S | D | G | T | F | T | S | E | L | S | R | L | A | E | G | A | R | L | Q | R | L | L | Q | G | L | V | 13,38 | 2,04 | 106,30 | 3,58 | | |
| 16 | H | S | D | G | T | F | T | S | E | L | S | R | L | R | A | G | A | R | L | Q | R | L | L | Q | G | L | V | 16,41 | 3,16 | 105,27 | 7,71 | | |
| 17 | H | S | D | G | T | F | T | S | E | L | S | R | L | R | E | A | A | R | L | Q | R | L | L | Q | G | L | V | 2,63 | 0,90 | 108,87 | 7,40 | | |
| 18 | H | S | D | G | T | F | T | S | E | L | S | R | L | R | E | G | A | A | L | Q | R | L | L | Q | G | L | V | 5,37 | 1,66 | 110,97 | 5,59 | | |
| 19 | H | S | D | G | T | F | T | S | E | L | S | R | L | R | E | G | A | R | A | Q | R | L | L | Q | G | L | V | 62,53 | 8,55 | 107,03 | 4,54 | | |
| 20 | H | S | D | G | T | F | T | S | E | L | S | R | L | R | E | G | A | R | L | A | R | L | L | Q | G | L | V | 4,60 | 3,31 | 108,60 | 13,30 | | |
| 21 | H | S | D | G | T | F | T | S | E | L | S | R | L | R | E | G | A | R | L | Q | A | L | L | Q | G | L | V | 6,35 | 1,75 | 113,40 | 5,01 | | |
| 22 | H | S | D | G | T | F | T | S | E | L | S | R | L | R | E | G | A | R | L | Q | R | A | L | Q | G | L | V | 23,86 | 4,34 | 110,87 | 4,31 | | |
| 23 | H | S | D | G | T | F | T | S | E | L | S | R | L | R | E | G | A | R | L | Q | R | L | A | Q | G | L | V | 20,76 | 3,75 | 108,53 | 4,58 | | |
| 24 | H | S | D | G | T | F | T | S | E | L | S | R | L | R | E | G | A | R | L | Q | R | L | L | A | G | L | V | 4,71 | 2,01 | 111,70 | 6,77 | | |
| 25 | H | S | D | G | T | F | T | S | E | L | S | R | L | R | E | G | A | R | L | Q | R | L | L | Q | A | L | V | 4,12 | 0,49 | 119,50 | 9,15 | | |
| 26 | H | S | D | G | T | F | T | S | E | L | S | R | L | R | E | G | A | R | L | Q | R | L | L | Q | G | A | V | 14,96 | 2,19 | 112,30 | 4,00 | | |
| 27 | H | S | D | G | T | F | T | S | E | L | S | R | L | R | E | G | A | R | L | Q | R | L | L | Q | G | L | A | 7,20 | 2,60 | 114,57 | 7,50 | | |
| 28 | R | S | D | G | T | F | T | S | E | L | S | R | L | R | E | G | A | R | L | Q | R | L | L | Q | G | L | V | 475,36 | 166,97 | 66,47 | 6,36 | | |
| 29 | E | S | D | G | T | F | T | S | E | L | S | R | L | R | E | G | A | R | L | Q | R | L | L | Q | G | L | V | 671,37 | 180,62 | 63,70 | 3,24 | | |
| 30 | F | S | D | G | T | F | T | S | E | L | S | R | L | R | E | G | A | R | L | Q | R | L | L | Q | G | L | V | 97,34 | 22,95 | 67,60 | 5,09 | | |
| 31 | H | W | D | G | T | F | T | S | E | L | S | R | L | R | E | G | A | R | L | Q | R | L | L | Q | G | L | V | 41,63 | 24,81 | 20,17 | 4,55 | | |
| 32 | H | R | D | G | T | F | T | S | E | L | S | R | L | R | E | G | A | R | L | Q | R | L | L | Q | G | L | V | - | - | - | - | | |
| 33 | H | D | D | G | T | F | T | S | E | L | S | R | L | R | E | G | A | R | L | Q | R | L | L | Q | G | L | V | - | - | - | - | | |
| 34 | H | S | R | G | T | F | T | S | E | L | S | R | L | R | E | G | A | R | L | Q | R | L | L | Q | G | L | V | - | - | - | - | | |
| 35 | H | S | N | G | T | F | T | S | E | L | S | R | L | R | E | G | A | R | L | Q | R | L | L | Q | G | L | V | 315,45 | 123,95 | 105,20 | 9,30 | | |
| 36 | H | S | W | G | T | F | T | S | E | L | S | R | L | R | E | G | A | R | L | Q | R | L | L | Q | G | L | V | - | - | - | - | | |
| 37 | H | S | D | R | T | F | T | S | E | L | S | R | L | R | E | G | A | R | L | Q | R | L | L | Q | G | L | V | - | - | - | - | | |
| 38 | H | S | D | E | T | F | T | S | E | L | S | R | L | R | E | G | A | R | L | Q | R | L | L | Q | G | L | V | - | - | - | - | | |
| 39 | H | S | D | I | T | F | T | S | E | L | S | R | L | R | E | G | A | R | L | Q | R | L | L | Q | G | L | V | - | - | - | - | | |
| 40 | H | S | D | G | R | F | T | S | E | L | S | R | L | R | E | G | A | R | L | Q | R | L | L | Q | G | L | V | 4,16 | 0,70 | 106,47 | 4,48 | | |
| 41 | H | S | D | G | D | F | T | S | E | L | S | R | L | R | E | G | A | R | L | Q | R | L | L | Q | G | L | V | 4,10 | 0,84 | 103,67 | 6,37 | | |
| 42 | H | S | D | G | C | F | T | S | E | L | S | R | L | R | E | G | A | R | L | Q | R | L | L | Q | G | L | V | 9,66 | 1,70 | 107,17 | 7,43 | | |
| 43 | H | S | D | G | T | R | T | S | E | L | S | R | L | R | E | G | A | R | L | Q | R | L | L | Q | G | L | V | - | - | - | - | | |
| 44 | H | S | D | G | T | D | T | S | E | L | S | R | L | R | E | G | A | R | L | Q | R | L | L | Q | G | L | V | - | - | - | - | | |
| 45 | H | S | D | G | T | V | T | S | E | L | S | R | L | R | E | G | A | R | L | Q | R | L | L | Q | G | L | V | 61,03 | 22,75 | 109,27 | 2,75 | | |
| 46 | H | S | D | G | T | F | R | S | E | L | S | R | L | R | E | G | A | R | L | Q | R | L | L | Q | G | L | V | - | - | - | - | | |
| 47 | H | S | D | G | T | F | D | S | E | L | S | R | L | R | E | G | A | R | L | Q | R | L | L | Q | G | L | V | - | - | - | - | | |
| 48 | H | S | D | G | T | F | C | S | E | L | S | R | L | R | E | G | A | R | L | Q | R | L | L | Q | G | L | V | 540,62 | 288,00 | 102,57 | 11,14 | | |
| 49 | H | S | D | G | T | F | T | R | E | L | S | R | L | R | E | G | A | R | L | Q | R | L | L | Q | G | L | V | - | - | - | - | | |
| 50 | H | S | D | G | T | F | T | D | E | L | S | R | L | R | E | G | A | R | L | Q | R | L | L | Q | G | L | V | 38,99 | 6,87 | 86,87 | 3,95 | | |
| 51 | H | S | D | G | T | F | T | W | E | L | S | R | L | R | E | G | A | R | L | Q | R | L | L | Q | G | L | V | - | - | - | - | | |
| 52 | H | S | D | G | T | F | T | S | R | L | S | R | L | R | E | G | A | R | L | Q | R | L | L | Q | G | L | V | 18,64 | 2,30 | 94,23 | 11,42 | | |
| 53 | H | S | D | G | T | F | T | S | L | L | S | R | L | R | E | G | A | R | L | Q | R | L | L | Q | G | L | V | 15,98 | 5,09 | 106,43 | 7,73 | | |
| 54 | H | S | D | G | T | F | T | S | Y | L | S | R | L | R | E | G | A | R | L | Q | R | L | L | Q | G | L | V | 11,75 | 1,29 | 103,23 | 7,20 | | |
| 55 | H | S | D | G | T | F | T | S | E | R | S | R | L | R | E | G | A | R | L | Q | R | L | L | Q | G | L | V | 154,10 | 58,03 | 86,62 | 7,70 | | |
| 56 | H | S | D | G | T | F | T | S | E | D | S | R | L | R | E | G | A | R | L | Q | R | L | L | Q | G | L | V | 869,29 | 640,61 | 62,80 | 12,45 | | |
| 57 | H | S | D | G | T | F | T | S | E | N | S | R | L | R | E | G | A | R | L | Q | R | L | L | Q | G | L | V | 213,65 | 121,19 | 83,67 | 11,03 | | |
| 58 | H | S | D | G | T | F | T | S | E | L | S | R | L | R | E | G | A | R | L | Q | R | L | L | R | G | L | V | 2,95 | 0,65 | 113,73 | 3,59 | | |
| 59 | H | S | D | G | T | F | T | S | E | L | S | R | L | R | E | G | A | R | L | Q | R | L | L | E | G | L | V | 3,47 | 0,19 | 108,90 | 6,08 | | |
| 60 | H | S | D | G | T | F | T | S | E | L | S | R | L | R | E | G | A | R | L | Q | R | L | L | L | G | L | V | 4,14 | 1,80 | 116,10 | 3,22 | | |
| 61 | H | S | D | G | T | F | T | S | E | L | S | R | L | R | E | G | A | R | L | Q | R | L | L | Q | R | L | V | 5,25 | 2,09 | 112,00 | 2,00 | | |
| 62 | H | S | D | G | T | F | T | S | E | L | S | R | L | R | E | G | A | R | L | Q | R | L | L | Q | E | L | V | 4,32 | 2,03 | 111,30 | 4,24 | | |
| 63 | H | S | D | G | T | F | T | S | E | L | S | R | L | R | E | G | A | R | L | Q | R | L | L | Q | I | L | V | 7,44 | 4,42 | 106,87 | 0,85 | | |
| 64 | H | S | D | G | T | F | T | S | E | L | S | R | L | R | E | G | A | R | L | Q | R | L | L | Q | C | L | V | 20,93 | 10,02 | 110,93 | 0,75 | | |
| 65 | H | S | D | G | T | F | T | S | E | L | S | R | L | R | E | G | A | R | L | Q | R | L | L | Q | G | R | V | 9,06 | 3,94 | 105,47 | 1,21 | | |
| 66 | H | S | D | G | T | F | T | S | E | L | S | R | L | R | E | G | A | R | L | Q | R | L | L | Q | G | D | V | 15,32 | 3,85 | 105,00 | 2,78 | | |
| 67 | H | S | D | G | T | F | T | S | E | L | S | R | L | R | E | G | A | R | L | Q | R | L | L | Q | G | F | V | 12,00 | 4,16 | 103,53 | 2,33 | | |

### Supplementary table 2

Pharmacological properties of secretin variants, part 2.

| peptide | sequence | | | | | | | | | | | | | | | | | | | | | | | | | | | β-arrestin2-GFP agonism | | | | cAMP agonism | |
|---|---|---|---|---|---|---|---|---|---|---|---|---|---|---|---|---|---|---|---|---|---|---|---|---|---|---|---|---|---|---|---|---|---|
| | 1 | 2 | 3 | 4 | 5 | 6 | 7 | 8 | 9 | 10 | 11 | 12 | 13 | 14 | 15 | 16 | 17 | 18 | 19 | 20 | 21 | 22 | 23 | 24 | 25 | 26 | 27 | potency [nM] | SD [nM] | efficacy [% Sec] | SD [% Sec] | potency [nM] | SD [nM] |
| 68 | H | S | D | G | T | F | T | S | E | L | S | R | L | K | E | G | A | R | L | Q | R | L | L | Q | G | L | V | 9,53 | 1,58 | 114,30 | 7,33 | | |
| 69 | H | S | D | G | T | F | T | S | E | L | S | R | L | R | D | G | A | R | L | Q | R | L | L | Q | G | L | V | 4,52 | 1,61 | 111,23 | 6,90 | | |
| 70 | H | S | D | G | T | F | T | S | E | L | S | R | L | R | E | G | E | R | L | Q | R | L | L | Q | G | L | V | 2,26 | 0,71 | 104,77 | 2,89 | | |
| 71 | H | S | D | G | T | F | T | S | E | L | S | R | L | R | E | G | A | R | L | E | R | L | L | Q | G | L | V | 2,28 | 0,54 | 100,73 | 2,80 | | |
| 72 | H | S | D | G | T | F | T | S | E | L | S | R | L | R | E | L | A | R | L | Q | R | L | L | Q | G | L | V | 3,32 | 0,60 | 104,60 | 4,52 | | |
| 73 | P | S | D | G | T | F | T | S | E | L | S | R | L | R | E | G | A | R | L | Q | R | L | L | Q | G | L | V | 39,08 | 12,83 | 96,57 | 6,31 | | |
| 74 | H | P | D | G | T | F | T | S | E | L | S | R | L | R | E | G | A | R | L | Q | R | L | L | Q | G | L | V | 3,37 | 1,99 | 106,23 | 2,44 | | |
| 75 | H | S | P | G | T | F | T | S | E | L | S | R | L | R | E | G | A | R | L | Q | R | L | L | Q | G | L | V | - | - | - | - | | |
| 76 | H | S | D | P | T | F | T | S | E | L | S | R | L | R | E | G | A | R | L | Q | R | L | L | Q | G | L | V | - | - | - | - | | |
| 77 | H | S | D | G | P | F | T | S | E | L | S | R | L | R | E | G | A | R | L | Q | R | L | L | Q | G | L | V | - | - | - | - | | |
| 78 | H | S | D | G | T | P | T | S | E | L | S | R | L | R | E | G | A | R | L | Q | R | L | L | Q | G | L | V | - | - | - | - | | |
| 79 | H | S | D | G | T | F | P | S | E | L | S | R | L | R | E | G | A | R | L | Q | R | L | L | Q | G | L | V | - | - | - | - | | |
| 80 | H | S | D | G | T | F | T | P | E | L | S | R | L | R | E | G | A | R | L | Q | R | L | L | Q | G | L | V | - | - | - | - | | |
| 81 | H | S | D | G | T | F | T | S | P | L | S | R | L | R | E | G | A | R | L | Q | R | L | L | Q | G | L | V | - | - | - | - | | |
| 82 | H | S | D | G | T | F | T | S | E | P | S | R | L | R | E | G | A | R | L | Q | R | L | L | Q | G | L | V | - | - | - | - | | |
| 83 | H | S | D | G | T | F | T | S | E | L | S | R | L | R | E | G | A | R | L | Q | R | L | L | Q | P | L | V | 22,13 | 10,01 | 103,17 | 4,12 | | |
| 84 | H | S | D | G | T | F | T | S | E | L | S | R | L | R | E | G | A | R | L | Q | R | L | L | Q | G | P | V | 72,64 | 22,88 | 95,03 | 7,74 | | |
| 85 | H | S | D | G | T | F | T | S | E | L | S | R | L | R | E | G | A | R | L | Q | R | L | L | Q | G | L | P | 5,19 | 2,62 | 100,47 | 11,91 | | |
| 86 | h | S | D | G | T | F | T | S | E | L | S | R | L | R | E | G | A | R | L | Q | R | L | L | Q | G | L | V | 13,59 | 4,52 | 107,20 | 1,67 | | |
| 87 | H | s | D | G | T | F | T | S | E | L | S | R | L | R | E | G | A | R | L | Q | R | L | L | Q | G | L | V | 45,06 | 21,00 | 91,57 | 6,20 | | |
| 88 | H | S | d | G | T | F | T | S | E | L | S | R | L | R | E | G | A | R | L | Q | R | L | L | Q | G | L | V | 135,95 | 67,03 | 89,07 | 1,87 | | |
| 89 | H | S | D | G | t | F | T | S | E | L | S | R | L | R | E | G | A | R | L | Q | R | L | L | Q | G | L | V | 308,43 | 31,06 | 102,97 | 5,57 | | |
| 90 | H | S | D | G | T | f | T | S | E | L | S | R | L | R | E | G | A | R | L | Q | R | L | L | Q | G | L | V | - | - | - | - | | |
| 91 | H | S | D | G | T | F | t | S | E | L | S | R | L | R | E | G | A | R | L | Q | R | L | L | Q | G | L | V | 1.537,76 | 534,38 | 98,33 | 5,69 | | |
| 92 | H | S | D | G | T | F | T | s | E | L | S | R | L | R | E | G | A | R | L | Q | R | L | L | Q | G | L | V | 1.873,16 | 538,08 | 90,37 | 6,84 | | |
| 93 | H | S | D | G | T | F | T | S | e | L | S | R | L | R | E | G | A | R | L | Q | R | L | L | Q | G | L | V | 20,35 | 6,53 | 100,40 | 5,89 | | |
| 94 | H | S | D | G | T | F | T | S | E | l | S | R | L | R | E | G | A | R | L | Q | R | L | L | Q | G | L | V | 20,99 | 5,47 | 94,17 | 6,05 | | |
| 95 | H | S | D | G | T | F | T | S | E | L | S | R | L | R | E | G | A | R | L | Q | R | L | L | Q | G | l | V | 8,48 | 3,66 | 101,90 | 4,71 | | |
| 96 | H | S | D | G | T | F | T | S | E | L | S | R | L | R | E | G | A | R | L | Q | R | L | L | Q | G | V | v | 4,73 | 2,07 | 99,97 | 3,64 | | |
| 97 | H | S | D | G | T | F | T | S | E | L | S | R | L | R | E | G | A | R | L | Q | R | L | L | Q | G | L | | 8,80 | 2,92 | 109,53 | 9,99 | 0,51 | 0,17 |
| 98 | H | S | D | G | T | F | T | S | E | L | S | R | L | R | E | G | A | R | L | Q | R | L | L | Q | G | | | 26,54 | 4,55 | 104,47 | 14,11 | 1,55 | 0,26 |
| 99 | H | S | D | G | T | F | T | S | E | L | S | R | L | R | E | G | A | R | L | Q | R | L | L | Q | | | | 35,92 | 7,43 | 103,60 | 9,27 | 2,09 | 0,43 |
| 100 | H | S | D | G | T | F | T | S | E | L | S | R | L | R | E | G | A | R | L | Q | R | L | L | | | | | 130,92 | 21,82 | 92,23 | 5,84 | 7,63 | 1,27 |
| 101 | H | S | D | G | T | F | T | S | E | L | S | R | L | R | E | G | A | R | L | Q | R | L | | | | | | | | | | 57,02 | 3,56 |
| 102 | H | S | D | G | T | F | T | S | E | L | S | R | L | R | E | G | A | R | L | Q | R | | | | | | | - | - | - | - | 585,95 | 531,03 |
| 103 | H | S | D | G | T | F | T | S | E | L | S | R | L | R | E | G | A | R | L | Q | | | | | | | | - | - | - | - | 874,80 | 718,23 |
| 104 | H | S | D | G | T | F | T | S | E | L | S | R | L | R | E | G | A | R | L | | | | | | | | | - | - | - | - | 2.213,08 | 1.474,81 |
| 105 | H | S | D | G | T | F | T | S | E | L | S | R | L | R | E | G | A | R | | | | | | | | | | - | - | - | - | 4.710,28 | 3.157,82 |
| 106 | H | S | D | G | T | F | T | S | E | L | S | R | L | R | E | G | A | | | | | | | | | | | - | - | - | - | 9.433,15 | 7.095,81 |
| 107 | H | S | D | G | T | F | T | S | E | L | S | R | L | R | E | G | | | | | | | | | | | | - | - | - | - | 20.152,03 | 14.941,03 |
| 108 | H | S | D | G | T | F | T | S | E | L | S | R | L | R | E | | | | | | | | | | | | | - | - | - | - | - | - |
| 109 | H | S | D | G | T | F | T | S | E | L | S | R | L | R | | | | | | | | | | | | | | - | - | - | - | 6.025,18 | 5.317,17 |
| 110 | H | S | D | G | T | F | T | S | E | L | S | R | L | | | | | | | | | | | | | | | - | - | - | - | 11.619,34 | |
| 111 | H | S | D | G | T | F | T | S | E | L | S | R | | | | | | | | | | | | | | | | - | - | - | - | - | - |
| 112 | H | S | D | G | T | F | T | S | E | L | S | | | | | | | | | | | | | | | | | - | - | - | - | - | - |
| 113 | H | S | D | G | T | F | T | S | E | L | | | | | | | | | | | | | | | | | | - | - | - | - | - | - |
| 114 | H | S | D | G | T | F | T | S | E | | | | | | | | | | | | | | | | | | | - | - | - | - | 12.600,00 | |
| 115 | | S | D | G | T | F | T | S | E | L | S | R | L | R | E | G | A | R | L | Q | R | L | L | Q | G | L | V | - | - | - | - | 21,96 | |
| 116 | | | D | G | T | F | T | S | E | L | S | R | L | R | E | G | A | R | L | Q | R | L | L | Q | G | L | V | - | - | - | - | 148,71 | |
| 117 | | | | G | T | F | T | S | E | L | S | R | L | R | E | G | A | R | L | Q | R | L | L | Q | G | L | V | - | - | - | - | 165,99 | |
| 118 | | | | | T | F | T | S | E | L | S | R | L | R | E | G | A | R | L | Q | R | L | L | Q | G | L | V | - | - | - | - | 578,32 | 359,87 |
| 119 | | | | | | F | T | S | E | L | S | R | L | R | E | G | A | R | L | Q | R | L | L | Q | G | L | V | - | - | - | - | 1.112,83 | 671,19 |
| 120 | | | | | | | T | S | E | L | S | R | L | R | E | G | A | R | L | Q | R | L | L | Q | G | L | V | - | - | - | - | 3.366,30 | 1.570,29 |
| 121 | | | | | | | | S | E | L | S | R | L | R | E | G | A | R | L | Q | R | L | L | Q | G | L | V | - | - | - | - | - | - |
| 122 | | | | | | | | | E | L | S | R | L | R | E | G | A | R | L | Q | R | L | L | Q | G | L | V | - | - | - | - | 5.852,93 | 762,96 |
| 123 | | | | | | | | | | L | S | R | L | R | E | G | A | R | L | Q | R | L | L | Q | G | L | V | - | - | - | - | 9.495,41 | 5.881,85 |
